# BioBatchNet: A Dual-Encoder Framework for Robust Batch Effect Correction in Imaging Mass Cytometry

**DOI:** 10.1101/2025.03.15.643447

**Authors:** Haiping Liu, Shaojie Zhang, Shengzhong Mao, Qian Zhao, Yuxi Zhou, Andrew Gilmore, Mauricio A. Alvarez, Hongpeng Zhou

## Abstract

**Motivation:** Imaging Mass Cytometry (IMC) is a cutting-edge technology for analysing spatially resolved protein expression at the single-cell level. However, its downstream analyses are often hindered by batch effects, which introduce systematic biases and obscure true biological variations. Existing correction methods, largely developed for scRNA-seq data, struggle to achieve precise control, leading to either over-correction by removing critical biological information, or under-correction by leaving residual batch effects. Moreover, these methods face challenges in adapting to IMC data due to differences in data characteristics. Furthermore, IMC data often feature imbalanced and overlapping cell populations, complicating clustering and downstream analysis. These challenges underscore the need for a robust and controllable batch effect correction approach tailored to IMC data.

**Results:** We present BioBatchNet, a dual-encoder framework utilising adversarial training to explicitly disentangle batch-specific and biological signals. BioBatchNet enables controllable batch effect correction, effectively balancing correction with the preservation of biological variation. We evaluated BioBatchNet on three IMC datasets, where it outperformed seven benchmarking methods with robustness in both correction and biological signal conservation. Additionally, we developed a Constrained Pairwise Clustering (CPC) method, which employs constrained pairs to improve clustering performance, even in datasets with imbalanced and overlapping cell populations. To validate its generalisability, BioBatchNet was also applied to four scRNA-seq datasets, where it delivered competitive performance compared to eight typical methods. These results demonstrate BioBatchNet’s generalisability and robustness in correcting batch effects across diverse single-cell datasets and underscore its potential for large-scale biological analyses.

## 1. Introduction

Imaging Mass Cytometry (IMC) is an advanced technology that enables simultaneous staining with multiple antibodies, allowing for the analysis of single-cell protein expression while preserving spatial context (Chang et al., 2017). These features make IMC a powerful tool for uncovering complex biological systems and disease mechanisms (Ali et al., 2020; Risom et al., 2022). However, downstream analyses of IMC data (e.g., clustering, cell type classification, etc.) are often hindered by batch effects. These systematic biases arise from technical or procedural inconsistencies (e.g., variations in experimental protocols, patient cohorts, etc.) during sample processing (Hunter et al., 2024; Tran et al., 2020; Shree et al., 2023). Existing batch effect correction methods, predominantly developed for scRNA-seq data, often fail to adapt to IMC data due to differences in data characteristics. Specifically, scRNA-seq data is characterized by discrete count values and comprehensive gene expression profiles (Lopez et al., 2018; Luecken et al., 2022; Peng et al., 2019), but IMC data involves continuous antibody intensity measurements and limited marker panels. Moreover, although correcting batch effects is essential, over-correction risks removing both batch effects and genuine biological signals, leading to misinterpretation. These challenges underscore the need for a robust batch effect correction method tailored to IMC data, which can effectively correct batch effects while preserving biological variation for accurate downstream analysis.

Existing batch effect correction methods, primarily developed for scRNA-seq data, can be broadly categorised into parametric, Mutual Nearest Neighbor (MNN)-based, and embedding-based approaches. Parametric method, such as ComBat, uses empirical Bayes shrinkage to achieve linear batch effect correction but cannot perform well for datasets with non-linear or complex batch effects (Johnson et al., 2007). MNN-based methods, including MNN (Haghverdi et al., 2018), Fast-MNN (Zhang et al., 2019), BBKNN (Polański et al., 2020), Harmony (Korsunsky et al., 2019), scDML (Yu et al., 2023) and Scanorama (Hie et al., 2019), correct batch effects by identifying and aligning similar data across batches. While effective for alignment, these methods face scalability issues with large-scale datasets (Tran et al., 2020; Li et al., 2023; Yu et al., 2023) and lack of control over the degree of correction, making it challenging to achieve a balance between batch effect removal and biological conservation. Moreover, MNN-based methods rely on similarity calculations, which are difficult to standardise across omics data with distinct distributions, limiting their generalisability.

Embedding-based methods, on the other hand, achieve batch effect correction by learning batch-corrected embedding through latent space models such as Generative Adversarial Networks (GANs), Autoencoders (AEs) and Variational Autoencoders (VAEs). A notable example is scVI (Lopez et al., 2018), which employs a VAE model to disentangle batch-unrelated information while using batch IDs to model batch effects. scVI has been subsequently adopted as the backbone for other scRNA-seq data analysis methods (Xu et al., 2021; Shree et al., 2023; Zhang et al., 2024; Xiong et al., 2024; Ashuach et al., 2023). However, due to its over-reliance on batch IDs to encode batch-specific information, scVI and its variants often fail to completely remove batch effects, leaving residual biases in the latent space and resulting in suboptimal performance. Alternatively, iMAP (Wang et al., 2021) uses GAN to disentangle biological features but adopting MNN for integration, limiting its scalability to large datasets. Recent advancements, such as scDC (Li et al., 2023) and AIF (Monnier and Cournède, 2024), have explored to use independent or shared encoders to disentangle biological and batch-specific information directly from raw data. However, scDC lacks precise control over the extent of correction and is constrained to clustering tasks, while AIF’s shared encoder tends to mix batch and biological signals, making effective disentanglement challenging.

Considering these challenges and the lack of studies specially addressing IMC data, we developed BioBatchNet, a dual-encoder VAE framework to tackle these issues. By integrating adversarial techniques, BioBatchNet can explicitly disentangle biological signals from batch-specific information. The structure of BioBatchNet comprises three core components: a Bio-encoder, a Batch-encoder, and a decoder. The Bio-encoder, guided by a batch discriminator (Ganin et al., 2016), extracts shared biological signals across batches, while the Batch-encoder learns batch-specific information from the data with the aid of a supervised batch classifier. These two encoders ensure that BioBatchNet can achieve effective disentanglement in an unsupervised manner. Finally, the decoder integrates the outputs from both encoders to reconstruct the original data. Besides, BioBatchNet enables precise control over the extent of batch effect correction by finetuning hyperparameters that regulate the strength of adversarial interactions. It should also be noted that while primarily designed for IMC data, BioBatchNet is also versatile by seamlessly adapting to scRNA-seq data, showcasing its potential for broader applications in single-cell analyses.

In addition to addressing batch effect correction, the limited marker information in IMC data often leads to overlapping cell populations, presenting significant challenges for clustering, a critical step in downstream analysis (Risom et al., 2022; Schapiro et al., 2017; Levine et al., 2015; Li et al., 2023; Yu et al., 2023). To tackle this issue, we developed a Constrained Pairwise Clustering (CPC) method that leverage prior knowledge to enhance clustering performance and can be integrated into BioBatchNet. CPC uses low-cost pairwise constraints (i.e., “must-link” (ML) and “cannot-link” (CL)) to guide the clustering process, where ML denotes samples belonging to the same cluster, and CL indicates samples from different clusters (Zhang et al., 2020a). Unlike traditional clustering methods, which often struggles in imbalanced and overlapping datasets (e.g., Leiden (Traag et al., 2019), KMeans, Phenograph (Levine et al., 2015), DEC (Xie et al., 2016), scDeepCluster (Tian et al., 2019), etc) or rely heavily on extensive annotations (Tian et al., 2021b; Cao et al., 2020), CPC employes weak supervision to significantly improve clustering reliability and interpretability. By integrating deep embedded clustering framework (DEC), scDCC (Tian et al., 2021a) is the first approach to apply constrained pairs to single-cell data. However, scDCC can not deal with imbalanced or overlapping data caused by the intrinsic limitations of DEC (Xie et al., 2016). In contrast, our CPC method eliminates the dependency on DEC and solely utilise constrained pairs as the weak prior information to guide the clustering process. Our method achieves a substantial improvement in clustering accuracy and reliability, making it a robust solution for the challenges posed by IMC data.

BioBatchNet is the first method to simultaneously achieve disentangled and controllable batch effect correction while incorporating constrained clustering specifically for IMC data. The key contributions of this work include:

- **Robust batch effect correction**: BioBatchNet leverages a novel dual-encoder VAE structure to effectively disentangle biological signals from batch effects, providing precise and controllable correction. Experimental results demonstrate that BioBatchNet consistently outperforms seven state-of-the-art benchmarking methods.
- **Improved clustering**: By integrating a Constrained Pairwise Clustering (CPC) method, BioBatchNet significantly enhances clustering accuracy using weak but cost-effective prior knowledge. This approach establishes a new standard for constrained clustering in IMC data, particularly for challenging scenarios with imbalanced and overlapping cell populations.
- **Generalisability**: BioBatchNet demonstrates versatility and scalability by delivering competitive performance across diverse datasets, including three IMC and four scRNA-seq datasets. The method achieves robust results in both batch effect correction and clustering tasks, highlighting its potential as a comprehensive solution for single-cell data analysis.

## 2. Methods

### 2.1. Model Structure

As illustrated in Figure 1, BioBatchNet comprises two independent encoders: (i) An encoder paired with a batch discriminator for encoding biological signals, referred to as the Bio-encoder; (ii) An encoder with an auxiliary batch classifier for modelling batch-specific information, referred to as the Batch-encoder.

**Fig. 1:**
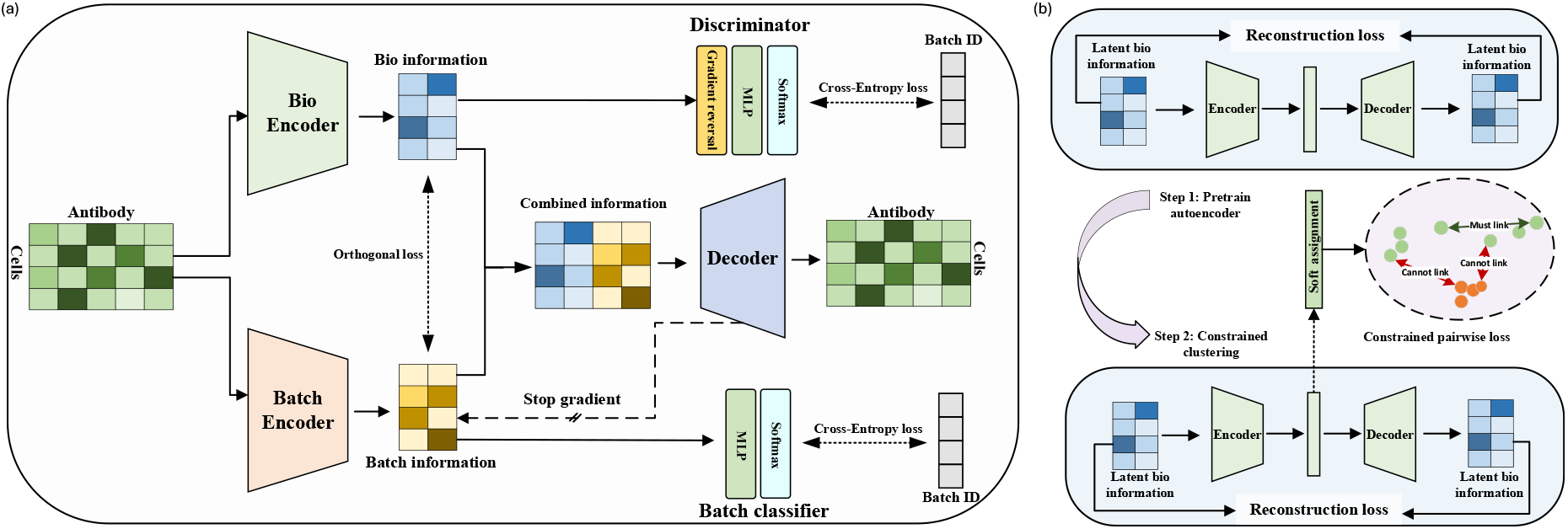
Overview of BioBatchNet framework and CPC optimisation workflow. (a) BioBatchNet is designed with a dual-encoder structure (i.e., Bio-encoder and Batch-encoder) to disentangle batch-specific and biological signals. The Bio-encoder is paired with a batch discriminator, optimised via adversarial training with a gradient reversal layer, to encode biological signals. The Batch-encoder is paired with a batch classifier to encode batch-specific information. (b) The CPC optimisation workflow consists of two steps. Step 1 involves pre-training an Autoencoder (AE), using the learned biological latent representation from BioBatchNet as input. Step 2 involves the refinement process that leverages pairwise constraints (i.e., must-link and cannot-link) to improve clustering accuracy.

Specifically, the Bio-encoder encodes the IMC data into biological latent representations which are fed into a batch discriminator equipped with a gradient reversal layer (Ganin et al., 2016) for adversarial training. With this adversarial setup, the Bio-encoder attempts to deceive the discriminator, while the discriminator strives to identify the batch source. Such adversarial interaction ensures that biological signals remain independent of batch effects. To compensate for the missing batch information in biological latent representations, which is critical for high-quality reconstruction, we introduce a batch encoder to learn batch-specific latent representations directly from the raw data. To ensure that these batch-specific latent representations exclusively capture batch-related information, they are fed into a batch classifier. Additionally, to prevent the batch encoder from being influenced by gradients related to the biological information, we block gradient back-propagation from the decoder and batch encoder. Furthermore, an orthogonal loss function is developed to enforce the independence between biological latent representations and batch-specific latent representations. Finally, these two types of latent representations are concatenated and used as input to the decoder for data reconstruction.

### 2.2. Optimisation Algorithm

In this section, we first derive the Evidence Lower Bound (ELBO) tailored to BioBatchNet, which forms the foundation of our loss function. Next, we illustrate our adversarial disentanglement strategy, which is designed to effectively correct batch effect while preserving biological signals.

#### 2.2.1. Evidence Lower Bound Derivation

Similar to other VAE models, BioBatchNet is optimised by maximising the ELBO, which provides a tractable lower bound to the marginal likelihood. Specifically, given the raw data *x*, the marginal likelihood can be expressed as:

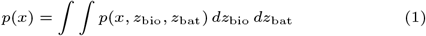

where *z*_bio_ and *z*_bat_ denote the biological and batch-specific latent representation, respectively. Since the integral in Eq. (1) is intractable, variational distributions *q*(*z*_bio_|*x*) and *q*(*z*_bat_|*x*) are introduced to approximate the true posteriors *p*(*z*_bio_|*x*) and *p*(*z*_bat_|*x*).

Using Jensen’s inequality (Zhang et al., 2018), the log-marginal likelihood can be lower-bounded as:

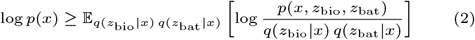

This is where the ELBO arises, providing a tractable objective for optimizing the model. Benefiting from our dual-encoders model structure, which disentangles *z*_bio_ and *z*_bat_ as independent vectors, the joint distribution *p*(*x, z*_bio_, *z*_bat_) can be further factorized as *p*(*x, z*_bio_, *z*_bat_) = *p*(*x*|*z*_bio_, *z*_bat_) *p*(*z*_bio_) *p*(*z*_bat_). Substituting this factorisation, the ELBO can be reformulated as:

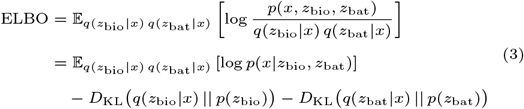

where the first term is used for data reconstruction, which is equivalent to Mean Squared Error (MSE) if we assume *p*(*x*|*z*_bio_, *z*_bat_) following aGuassian distribution. The second and third terms are the Kullback-Leibler divergences between the approximated posterior distributions and their respective priors *p*(*z*_bio_) and *p*(*z*_bat_). These priors are modelled as standard Gaussian distributions 𝒩 (0, 1) to ensure a well-regularised latent space.

It can be observed that the optimisation of ELBO enables an integration between reconstruction accuracy and the disentanglement of biological and batch-specific information. The disentanglement is further strengthened by our proposed disentangled representation learning strategy, which is detailed in the next section.

#### 2.2.2. Disentangled Representation Learning Strategy

To achieve disentanglement between biological and batch-specific information, we propose a disentangled representation learning strategy by employing three key techniques: adversarial training, gradient blocking, and incorporation of an orthogonal loss.

Adversarial training is implemented through two processes. The first focusses on the adversarial interaction between the bioencoder *E*_bio_ and the batch discriminator *D*_bat_. The role of *D*_bat_ is to identify and penalise any batch-related information encoded within the biological latent representation *z*_bio_ generated by *E*_bio_. Simultaneously, *E*_bio_ is trained to learn *z*_bio_ that excludes batch-related information, countering the efforts of *D*_bat_. This process is facilitated by a gradient reversal layer, leading to a min-max optimisation target as follows:

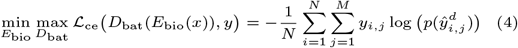

where *L*_ce_ is the cross-entropy loss, *y* denotes the batch labels, *N* is the number of samples, and *M* is the number of batches. 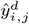 denotes the predicted output of *D*_bat_, with 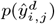 representing the probability of sample *i* being classfied into batch *j. y*_*i,j*_ denotes the true probability of sample *i* belonging to batch *j*.

The second process lies in the optimising dual encoder pathways with opposite training objective. As explained above, *E*_bio_ and *D*_bat_ are trained to prevent *z*_*bio*_ from including batch-related information. In contrast, the batch encoder *E*_bat_ and its classifier *C*_bat_ are trained by maximising classification accuracy to ensure *z*_*bat*_ captures as much batch-related information as possible. This adversarial dynamic strengthens the disentanglement of latent representations. The optimisation target for *E*_bat_ and *C*_bat_ can be defined as:

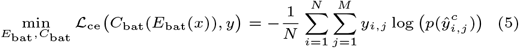

where 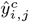 is the predicted output of *C*_bat_ and 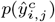 denotes the probability of sample *i* belonging to batch *j*.

To further ensure disentanglement, we employ gradient blocking to prevent the batch encoder from inadvertently capturing biological information. Specifically, gradients from the decoder to *E*_bat_ are blocked, ensuring *z*_bat_ remains purely batch-specific. Additionally, to enforce independence between *z*_bio_ and *z*_bat_, we introduce an orthogonal loss to penalize any linear correlation between them. First, the latent representations are centered by subtracting their respective means across batches:

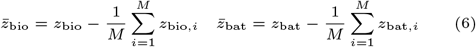

Then, the covariance matrix **C** between 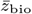 and 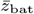 is computed:

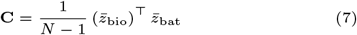

Finally, the orthogonal loss is defined as the squared Frobenius norm of **C**:

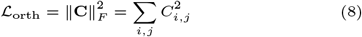

In conclusion, the overall loss function of BioBatchNet integrates the ELBO from Eq. (3), adversarial loss from Eq. (4), classification loss from Eq. (5), and the orthogonal loss from Eq. (8). To simplify notation, let *ℒ*_ce_ (*E*_bio_, *D*_bat_), *ℒ*_ce_ (*E*_bat_, *C*_bat_) denote the loss terms in Eq. (4) and Eq. (5), respectively. The overall loss function can be formulated as follows:

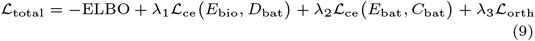

where *λ*_1_, *λ*_2_, *λ*_3_ are weighting factors that balance reconstruction accuracy and disentanglement objectives. And *λ*_1_ is the critical parameter controlling the extent of batch effect correction.

### 2.3. Constrained Pairwise Clustering Method

Constrained Pairwise Clustering (CPC) methods leverage weak priors, such as must-link (ML) and cannot-link (CL) constraints, to improve clustering reliability. Following prior work (Zhang et al., 2020a), we develop a two-step CPC approach that pretrains an Autoencoder (AE) and then refines the AE using pairwise constraints to optimise clustering, as illustrated in Figure 1(b).

The input to the AE is the latent biological representation *z*_bio_ generated by the Bio-encoder. The AE is pretrained with a standard MSE loss, enabling it encodes *z*_bio_ into a lower-dimensional representation suitable for clustering. After pretraining, a subset of data points is sampled and ML and CL pairs are defined within this subset. Soft assignment is then applied to assign samples to clusters based on the Student’s t-distribution. The clustering assignment probability *q*_*ij*_, denoting the probability of sample *i* belonging to cluster *j*, is defined as:

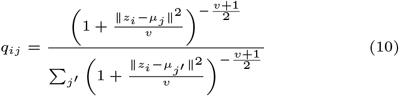

where *z*_*i*_ represents the latent representation of sample *i, µ*_*j*_ is the center of cluster *j*, and *v* denotes the degree of freedom in Student’s t-distribution.

To optimise the parameters in Eq. (10), the pairwise constrained loss are then incorporated to guide the learning process. Specifically, the loss for ML constraints set, which aims to enhance probability of samples in ML pairs being classified into the same cluster, can be formulated as:

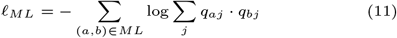

where (*a, b*) denotes the selected ML pair and *q*_*aj*_ and *q*_*bj*_ denote the probabilities of samples *a* and *b* belonging to cluster *j*. Similarly, the loss for CL constraints set, which aims to reduce probability of samples in CL pairs being classified into the same cluster, is defined as

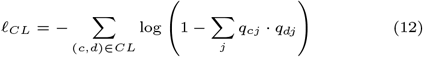

where (*c, d*) denotes the selected CL pair.

The total loss for constrained clustering combines reconstruction loss with the pairwise losses:

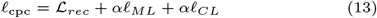

where ℒ _*rec*_ denotes the MSE reconstruction loss, *α* and *β* are weighting factors balancing the contributions of ML and CL losses. It should be note that, unlike previous CPC-based clustering method such as scDCC (Tian et al., 2021a), which replys on deep embedding clustering (DEC) framework (Xie et al., 2016) prone to poor performance on imbalanced and overlapping data (see supplementary material), our CPC approach discard DEC framework and directly utilises pairwise loss to update the soft assignment probability in Eq. (10), yielding a significant improvement in clustering reliability.

## 3. Experimental Results

### 3.1. Experimental Setup

#### 3.1.1. Datasets

We evaluated the effectiveness of BioBatchNet in three representative IMC datasets. The IMMUcan_Cancer dataset, collected from four cancer patients with distinct cancer indications, exhibits pronounced batch effects caused by patients’ variations (Windhager et al., 2023). The Damond Pancreas dataset focuses on type 1 diabetes, providing a large-scale benchmark for assessing scalability of our method (Damond et al., 2019). Finally, the Hoch_Melanoma dataset explores immune cell infiltration in melanoma samples and includes 39 batches, highlighting the necessity of batch effect correction with substantial batch variability (Hoch et al., 2022). To further validate the generalisation ability of BioBatchNet, we applied it to four scRNA-seq datasets, i.e., Macaque retina data obtained from (Peng et al., 2019), Human pancreas, Human immune, collected from Luecken et al. (2022) and Mouse brain data provided by Saunders et al. (2018); Rosenberg et al. (2018). These datasets span different tissues and species, offering diverse cell types and varying batch characteristics.

For data preprocessing, we applied standard preprocessing steps for scRNA-seq datasets, including the selection of highly variable genes, count normalization, and log1p transformation, except for scVI, which omits the log1p transformation step. In contrast, IMC datasets were used in their raw form without preprocessing.The details of all datasets (e.g., number of cells, number of batches, etc) and links to the data resources are provided in Table 1.

**Table 1.**
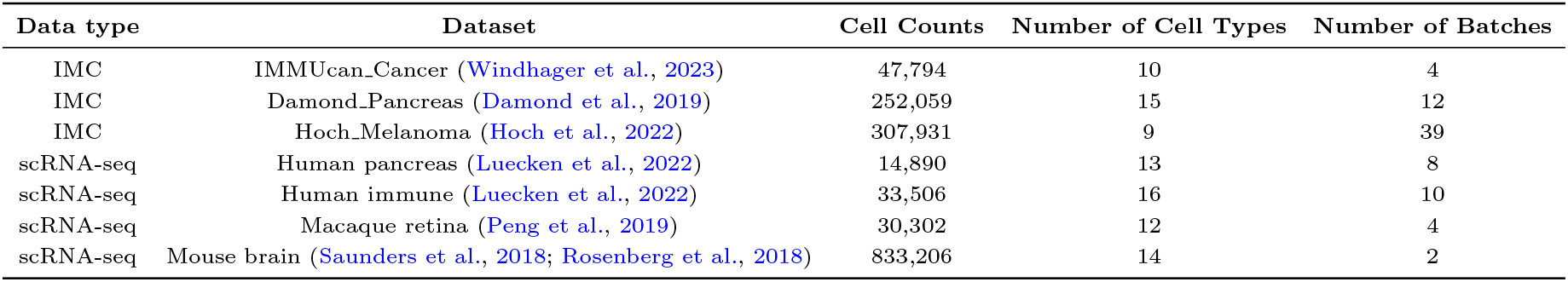
Summary of datasets used for evaluating BioBatchNet

| Data type | Dataset | Cell Counts | Number of Cell Types | Number of Batches |
| --- | --- | --- | --- | --- |
| IMC | IMMUCan.Cancer (Windhager et al., 2023) | 47,794 | 10 | 4 |
| IMC | Damond.Pancreas (Damond et al., 2019) | 252,059 | 15 | 12 |
| IMC | Hoch.Melanoma (Hoch et al., 2022) | 307,931 | 9 | 39 |
| scRNA-seq | Human pancreas (Luecken et al., 2022) | 14,890 | 13 | 8 |
| scRNA-seq | Human immune (Luecken et al., 2022) | 33,506 | 16 | 10 |
| scRNA-seq | Macaque retina (Peng et al., 2019) | 30,302 | 12 | 4 |
| scRNA-seq | Mouse brain (Saunders et al., 2018; Rosenberg et al., 2018) | 833,206 | 14 | 2 |

#### 3.1.2. Benchmarking Methods and Evaluation Metrics

To evaluate the performance of BioBatchNet on IMC datasets, we compared it against seven widely used batch effect correction methods: scVI (Lopez et al., 2018), Harmony (Korsunsky et al., 2019), BBKNN (Polański et al., 2020), Scanorama (Hie et al., 2019), Combat (Zhang et al., 2020b), iMAP (Wang et al., 2021), and scDREAMER (Shree et al., 2023). Notably, scDML (Yu et al., 2023) was excluded as it cannot process continuous data (Yu et al., 2023). To adapt scVI and scDREAMER for IMC datasets, we modified their output likelihood to Gaussian distribution,replacing their original Zero-Inflated Negative Binomial (ZINB) distribution designed for scRNA-seq datastes. For scRNA-seq datasets, we included scDML alongside above seven benchmark methods for comparison.

We evaluated BioBatchNet and benchmarking methods based on their performance in both batch effect correction and biological conservation. The performance in batch effect correction was evaluatedusing four metrics, i.e., iLISI, Graph-Connectivity, ASW-batch, PCR (see details in Section.B.1 of the Supplementary). Higher iLISI and Graph Connectivity values indicate better mixing of batches, whereas lower ASW-batch and PCR scores reflect reduced batch effects. The average of these metrics denotesthe batch effect correction score. Biological conservation was assessed using three metrics, i.e., ASW-cell, ARI and NMI (see details in Section.B.2 of Supplementary). Higher values for all three metrics indicate better preservation of biological variation, and their average represents the biological conservation score. Furthermore, to evaluate our developed constrained pairwise clustering method, we compared the CPC approach with K-means, Leiden algorithm, and scDCC (Tian et al., 2021a). Clustering performance was assessed using ARI, NMI, and ACC (see details in Section.B.3 of Supplementary). All metrics, including those for batch effect correction, biological conservation, and clustering performance, were normalised to a 0−1 scale using the Python scib package (Luecken et al., 2022).

### 3.2. Performance in IMC Datasets

In this subsection, experimental results of BioBatchNet on IMC datasets will be discussed. To validate the robustness of our method, repeated experiments with different random seed were implemented for each dataset. The best performance achieved by each method is summarized in Table 2. The dual-encoders, discriminator, classifier, and decoder in BioBatchNet are constructed using fully connected layers. Due to the page limitation, we present the result of IMMUcan_Cancer dataset as a representative example. Comprehensive results for other IMC datasets are included in Supplementary Materials (Section.C).

**Table 2.** Comparison of batch effect correction and biological conservation scores across batch correction methods in IMC datasets.

| Model \ Dataset | Batch Effect Correction Score |  |  | Biological Conservation Score |  |  |
| --- | --- | --- | --- | --- | --- | --- |
|  | IMMUcan_Cancer | Damond_Pancreas | Hoch_Melanoma | IMMUcan_Cancer | Damond_Pancreas | Hoch_Melanoma |
| scVI (Lopez et al., 2018) | 0.65 | 0.71 | 0.53 | 0.44 | 0.56 | 0.50 |
| Harmony (Korsunsky et al., 2019) | 0.82 | **0.80** | 0.67 | 0.53 | **0.56** | 0.53 |
| BBKNN (Polanski et al., 2020) | 0.56 | 0.61 | 0.48 | 0.39 | 0.41 | 0.52 |
| Scanorama (Hie et al., 2019) | 0.77 | 0.61 | 0.61 | 0.37 | 0.51 | 0.21 |
| Combat (Zhang et al., 2020b) | 0.78 | 0.73 | **0.75** | 0.51 | 0.57 | 0.48 |
| iMAP (Wang et al., 2021) | 0.72 | 0.48 | 0.48 | 0.36 | 0.29 | 0.26 |
| scDREAMER (Shree et al., 2023) | 0.83 | 0.75 | 0.71 | 0.39 | 0.35 | 0.36 |
| **BioBatchNet** | **0.87** | 0.77 | 0.69 | **0.56** | 0.56 | **0.58** |

The IMMUcan_Cancer dataset includes IMC samples from four patients with distinct cancer types, posing significant challenges due to the inherent strong batch effects, as evidenced by the clear separation of batches in the raw data (Figure 2.(a)). Figure 2.(a) and Figure 2.(b) visualise the latent biological representations generated by the Bio-encoder, which can illustrate the performance of batch effect correction and biological information conservation. It can be observed that Harmony, scDREAMER, and BioBatchNet effectively reduced batch effect with a better mixing of diverse batches (Figure 2.(a)). However, scDREAMER fails to preserve biological signals, as indicated by the lack of clear separation between cell types (Figure 2.(b)). In contrast, BioBatchNet sucessfully balanced batch effect correction with biological conservation, achieving the highest scores for both metrics (Figure 2.(d))

**Fig. 2:**
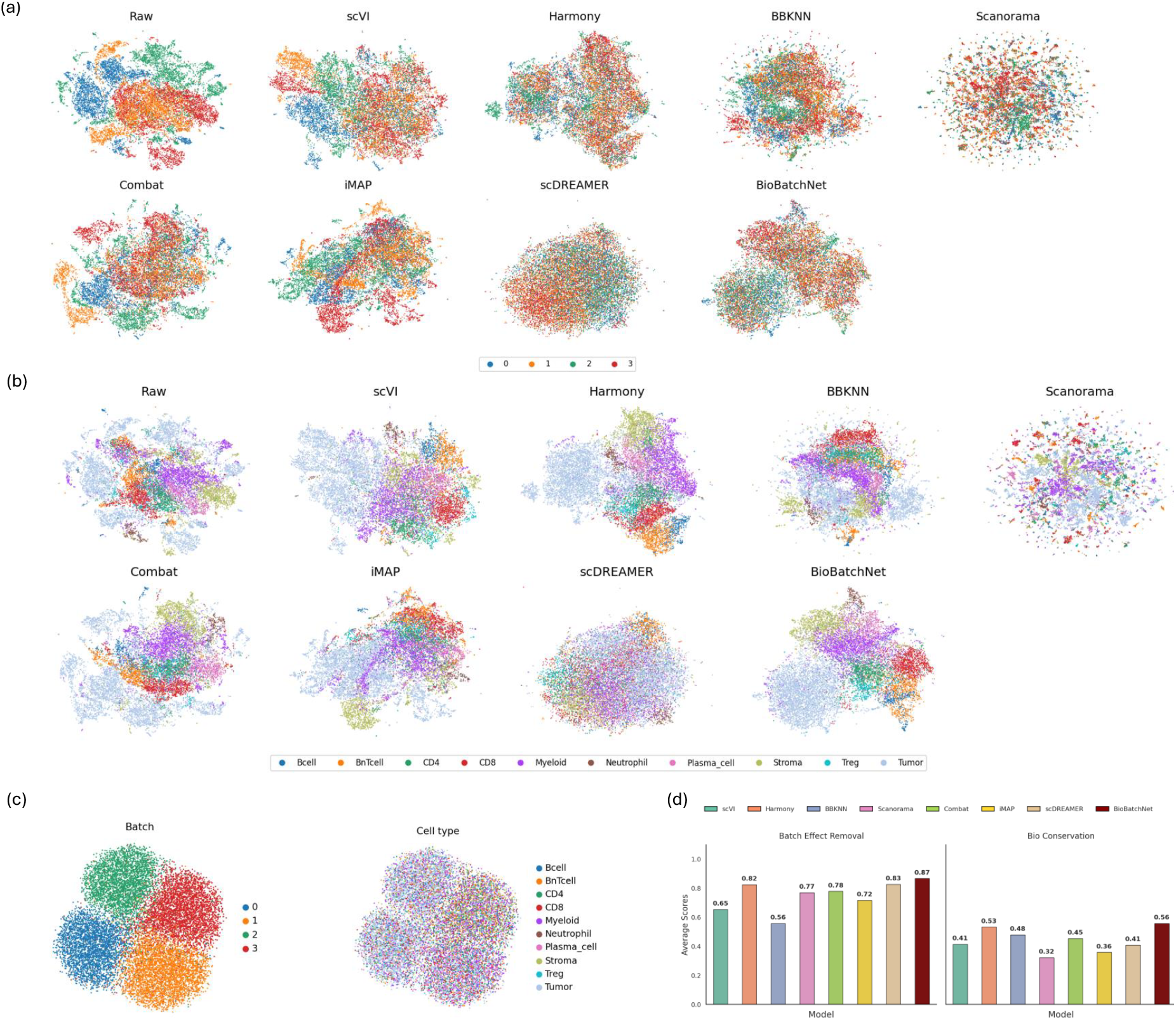
Comparison of methods for IMMUcan_Cancer dataset. **(a)** UMAP visualization illustrating of batch effect correction in latent biological representations. **(b)** UMAP visualization of biological signal conservation in latent biological representations. **(c)** UMAP visualization of batch-specific information encoded in latent batch-specific representations. **(d)** Quantitative comparison of batch effect correction and biological conservation scores.

In addition, IMC data often exhibit limited antibody diversity, leading to overlapping and closely related cell populations. In these cases, retaining some degree of overlap between biologically similar cell types is reasonable and expected (e.g., CD4 and CD8 T cells in the IMMUcan_Cancer dataset). Figure 2.(b) shows that BioBatchNet effectively preserved biological structures by maintaining overlaps between similar cell types while clearly separating distinct populations. Additionally, Figure 2.(c) illustrates the latent batch-specific representations generated by the Batch-encoder, highlighting a clear separation of batches without retaining biological signals, demonstrating BioBatchNet’s robust disentanglement capabilities.

On other IMC datasets, BioBatchNet maintained competitive preformance. Notably, BioBatchNet and Harmony can achieve similar result in the Damond_Pancreas dataset, which contains a large number of cells (see Supplementary Figure S1, Table S2). However, in the Hoch_Melanoma dataset, which features a high number of batches (39 batches), Harmony’s performance declines, particularly in the clustering accuracy, as reflected by a lower ARI value. In contrast, BioBatchNet excelled by achieving superior biological signal preservation while maintaining competitive batch effect correction (see Supplementary Figure S2 and Table S3). Other benchmarking methods cannot achieve the balance, often perform well on one aspect at the cost of the other. For example, Combat achieved the best batch effect correction performance with the cost of losing biological information. Overall, BioBatchNet demonstrated its ability to balance and optimise the batch effect correction and biological conservation across diverse IMC datasets. A more complete quantitative comparison for all methods is in Table 2.

### 3.3 Precision Control of Batch Effect Correction

Distinguishing batch effects from biological signals is inherently challenging, and the optimal level of correction often varies depending on the specific application. Therefore, a flexible method that allows precise adjustment of the correction degree is essential to avoid either over-correction or under-correction. Our BioBatchNet can enable controlling the degree of correction through modifying the weighting factor *λ*_1_ in the loss function, as defined in Eq. (9). A larger *λ*_1_ enforces a stronger correction.

To illustrate this, we conducted experiments to evaluate precision control on IMMUcan_Cancer dataset. As shown in Figure 3, when *λ*_1_ = 0.1, batch effects were insufficiently corrected, resulting in distinct clusters of tumor cells. Conversely, a higher *λ*_1_ = 0.7 led to over-correction causing a loss of biological information. An intermediate value of *λ*_1_ = 0.3 achieved an optimal trade-off, effectively mitigating batch effects while preserving critical biological information. The capability of precision control over batch effect correction enables BioBatchNet to adapt flexibly to diverse datasets with varying requirements for correction.

**Fig. 3:**
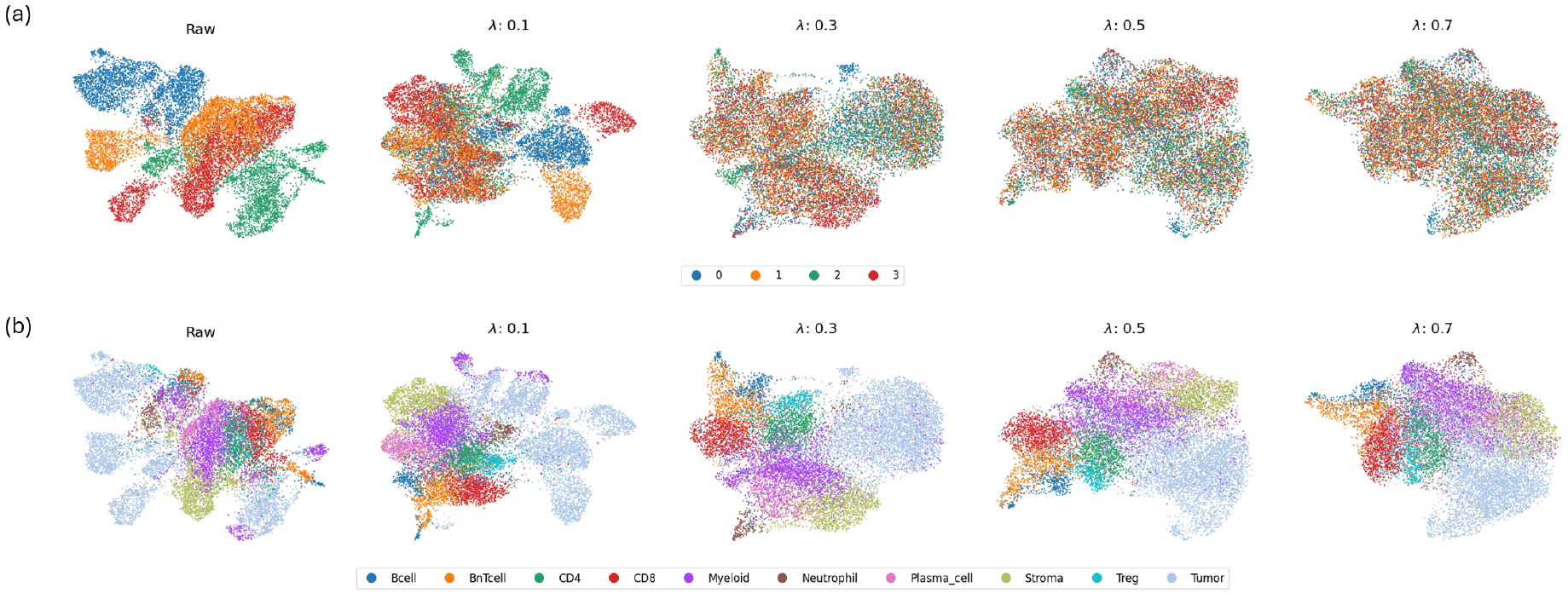
Precision control of batch effect correction on IMMUcan_Cancer dataset. (a) UMAP visualization showing the impact of weighting factor *λ*_1_ on batch effect correction. (b) UMAP visualization showing the impact of weighting factor *λ*_1_ on biological signal conservation.

### 3.4. Clustering Performance of CPC in IMC Datasets

As described in Section 2.3, we develop a Constrained Pairwise Clustering (CPC) method that leverages weak priors (i.e., ML and CL constraints) to enhance clustering accuracy and reliability. In our experiments, we defined 3, 000 ML or CL cell pairs as the weak prior. It should be noted that this approach to obtaining prior knowledge is practical and significantly cost-effective compared to acquiring strong prior knowledge, such as full cell-type annotations. For instance, in the Hoch_Melanoma dataset, these 3, 000 pairs accounts only a tiny fraction (approximately 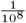) of all possible pairs.

Figure 4 illustrates how the inclusion of constrained pairs enhances clustering performance in IMMUcan_Cancer dataset. The result shows that the enhanced separation between CL pairs contributes to better separation of distinct clusters, while ML pairs improve alignment of similar clusters. Additional clustering results of CPC for Damond Pancreas and Hoch_Melanoma datasets can be found in Supplementary Section E.1. Table 3 ompares the clustering performance metrics of CPC with other methods, including K-means, Leiden, and scDCC. The CPC method consistently achieved the highest scores across all evaluation metrics. Notablly, scDCC, which is also a CPC-based method, failed to achieve competitive performance. This limitation arises because scDCC relies on the DEC framework (Xie et al., 2016), which struggles with imbalanced and overlapping cell populations, frequent challenges in IMC datasets (A detailed discussion on DEC is in Supplementary Section E.2). In contrast, our CPC approach avoids these issues by solely utilising pairwise loss to guide clustering, ensures more reliable and accurate results.

**Table 3.**
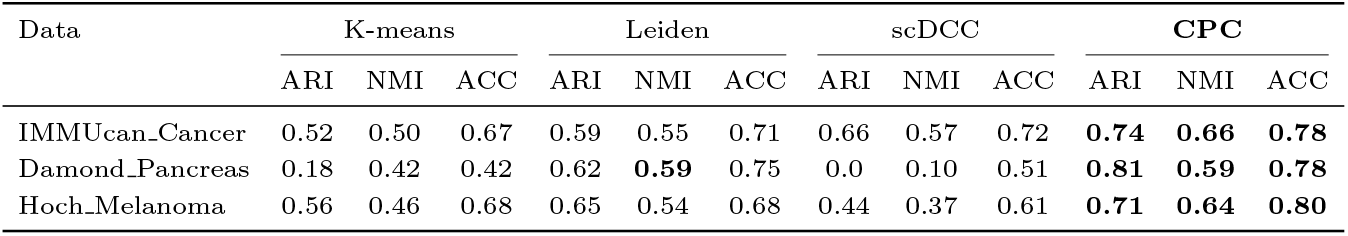
Comparison of clustering performance metrics across different methods in IMC datasets

| Data | K-means |  |  | Leiden |  |  | scDCC |  |  | CPC |  |  |
| --- | --- | --- | --- | --- | --- | --- | --- | --- | --- | --- | --- | --- |
|  | ARI | NMI | ACC | ARI | NMI | ACC | ARI | NMI | ACC | ARI | NMI | ACC |
| IMMUcan_Cancer | 0.52 | 0.50 | 0.67 | 0.59 | 0.55 | 0.71 | 0.66 | 0.57 | 0.72 | **0.74** | **0.66** | **0.78** |
| Damond_Pancreas | 0.18 | 0.42 | 0.42 | 0.62 | **0.59** | 0.75 | 0.0 | 0.10 | 0.51 | **0.81** | **0.59** | **0.78** |
| Hoch_Melanoma | 0.56 | 0.46 | 0.68 | 0.65 | 0.54 | 0.68 | 0.44 | 0.37 | 0.61 | **0.71** | **0.64** | **0.80** |

**Fig. 4:**
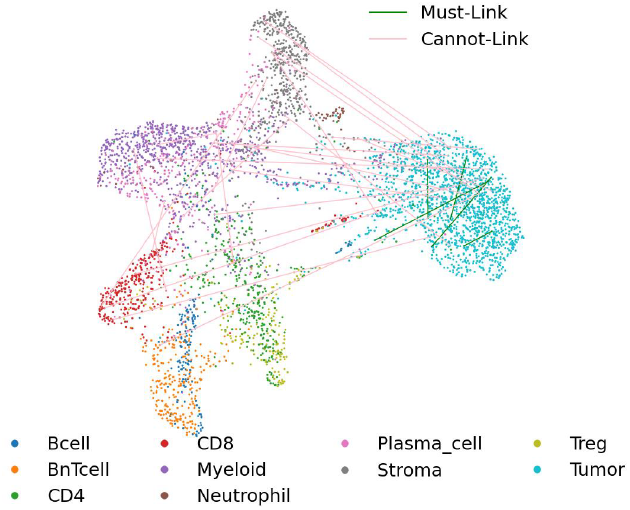
Improved clustering performance with the developed Constrained Pairwise Clustering approach in IMMUcan_Cancer dataset.

### 3.5. Generalization to scRNA-seq Datasets

To verify the generalisation ability of BioBatchNet, we tested it on four scRNA-seq datasets, i.e., Human pancreas, Human immune, Macaque retina, and Mouse brain. As shown in Table 4, BioBatchNet demonstrated competitive performance and robustness in balancing batch effect correction and biological conservation across these datasets. Specifically, for the Macaque retina (see Supplementary Figure S5) and Human pancreas datasets (see Supplementary Figure S3), BioBatchNet achieved the best batch effect removal performance, while maintaining competitive scores in biological conservation. In the Human immune dataset, which features overlapping cell populations similar as IMC datasets, BioBatchNet delivered competitive biological conservation scores and achieved the highest clustering performance metrics (i.e., ARI and NMI), although its batch effect correction performance was not the best (see Supplementary Table S5). For the Mouse brain dataset, which pose challenges due to the inherent noise in neural cells (see Supplementary Figure S6.(a)), BioBatchNet stood out as the only method capable of effectively removing batch effects while addressing the inherent noise, as demonstrated in Figure S6.(b) of the Supplementary Materials. It is also worth noting that to adapt BioBatchNet for scRNA-seq datasets, a tailored loss function was developed to address the unique characteristics of scRNA-seq data. The detailed mathematical derivation is provided in Section A.

**Table 4.** Comparison of batch effect correction and biological conservation scores across batch correction methods in scRNA-seq datasets

| Dataset<br>Model | Batch Effect Correction Score |  |  |  | Biological Conservation Score |  |  |  |
| --- | --- | --- | --- | --- | --- | --- | --- | --- |
|  | Human Pancreas | Immune | Macaque | Mouse Brain | Human Pancreas | Immune | Macaque | Mouse Brain |
| scVI (Lopez et al., 2018) | 0.67 | 0.71 | 0.77 | 0.67 | 0.83 | 0.65 | 0.80 | 0.47 |
| Harmony (Korsunsky et al., 2019) | 0.76 | 0.75 | 0.84 | 0.63 | 0.84 | 0.66 | 0.69 | 0.49 |
| BBKNN (Polański et al., 2020) | 0.46 | 0.50 | 0.47 | 0.38 | 0.82 | 0.61 | 0.76 | 0.27 |
| Scanorama (Hie et al., 2019) | 0.63 | 0.56 | 0.55 | 0.44 | 0.49 | 0.50 | 0.59 | 0.47 |
| Combat (Zhang et al., 2020b) | 0.77 | **0.80** | 0.79 | 0.68 | 0.77 | 0.68 | 0.69 | 0.48 |
| iMAP (Wang et al., 2021) | 0.76 | 0.74 | 0.69 | 0.70 | 0.80 | 0.61 | 0.62 | 0.49 |
| scDREAMER (Shree et al., 2023) | 0.78 | 0.67 | 0.81 | 0.70 | 0.87 | **0.72** | 0.80 | 0.51 |
| scDML (Yu et al., 2023) | 0.78 | 0.72 | 0.83 | 0.67 | **0.88** | 0.56 | **0.89** | 0.63 |
| BioBatchNet | **0.83** | 0.72 | **0.88** | **0.78** | 0.83 | 0.69 | 0.83 | **0.71** |

## 4. Conclusion

In this work, we present BioBatchNet, a novel dual-encoder VAE framework that combines adversarial techniques to disentangle biological signals and batch effect. BioBatchNet explicitly separates biological and batch-specific signals while enabling precise control over batch effect correction. Additionally, we integrate constrained clustering into our framework to enhance the interpretability and accuracy of downstream analyses. Through extensive evaluations on IMC and scRNA-seq datasets, we demonstrate that BioBatchNet outperforms state-of-the-art methods in both batch effect correction and clustering tasks, establishing a robust and scalable approach for single-cell data analysis.

In future work, we plan to explore the integration of spatial information into BioBatchNet, aiming to uncover deeper biological insights beyond clustering. One potential solution is to combine BioBatchNet’s latent embeddings with spatial data as input for Graph Neural Networks. This integration could provide a more comprehensive understanding of biological relationships and patterns. Additionally, we plan to explore other potential applications of BioBatchNet beyond batch effect correction, such as data imputation and data generation. Expanding its functionality in these areas could significantly enhance its utility and further demonstrate the versatility and effectiveness of our method.

## 5. Supplementary Data

Supplementary data is provided.

