## Supplementary material for "BioBatchNet: A Dual-Encoder Framework for Robust Batch Effect Correction in Imaging Mass Cytometry"

In the supplementary document, we first provide a detailed explanation of how we extend BioBatchNet to scRNA-seq data, along with its corresponding loss function in the section A. We then present the experimental results for both IMC in section C and scRNA-seq datasets in section D. Furthermore, we showcase the CPC results for the Damond and HochSchulz IMC datasets in section E.1. Lastly, we discuss the limitations of DEC in section E.2, which led us to exclude DEC and instead rely solely on constrained pairs for clustering.

#### A. Loss function of scRNA-seq data

##### A.1. Evidence Lower Bound

For scRNA-seq data, we start with the log-likelihood which should be maximized. As a result, given the input data $x$, we use not only the dual encoder framework to model the biological and batch latent representation $z_{bio}$ and $z_{batch}$, but also the library size $l$ as a scaling factor. Consequently, the log-likelihood can be written as:

$$\log p(x) = \log \int\int\int p(x, z_{\text{bio}}, z_{\text{bat}}, z_{\text{l}})\, dz_{\text{bio}}\, dz_{\text{bat}}\, dz_{\text{l}} \tag{14}$$

Since directly computing the above integral is intractable, we use variational inference to estimate the posterior distribution as follows:

$$\begin{aligned} \log p(x) &= \log \int\int\int p(x, z_{\text{bio}}, z_{\text{bat}}, z_{\text{l}})\, dz_{\text{bio}}\, dz_{\text{bat}}\, dz_{\text{l}} \\ &= \log \int\int\int \frac{p(x, z_{\text{bio}}, z_{\text{bat}}, z_{\text{l}})}{q(z_{\text{bio}}, z_{\text{bat}}, l|x)} q(z_{\text{bio}}, z_{\text{bat}}, l|x)\, dz_{\text{bio}}\, dz_{\text{bat}}\, dz_{\text{l}} \end{aligned} \tag{15}$$

where $z_{\text{bio}}$ and $z_{\text{bat}}$ are truly independent due to our dual-encoder framework and disentanglement strategy and $l$ is modeled under a mean-field assumption for simplicity. Thus, the variational poseterior can be factorized:

$$q(z_{\text{bio}}, z_{\text{bat}}, l|x) = q(z_{\text{bio}} \mid x)\, q(z_{\text{bat}} \mid x)\, q(l \mid x) \tag{16}$$

So the log-likelihood can be rewritten as

$$\log \mathbb{E}_{q_{\text{bio}}(z_{\text{bio}}|x)\, q_{\text{bat}}(z_{\text{bat}}\, q_{\text{l}}(z_{\text{l}})|x)} \left[ \frac{p(x, z_{\text{bio}}, z_{\text{bat}})}{q_{\text{bio}}(z_{\text{bio}}|x)\, q_{\text{bat}}(z_{\text{bat}}|x)\, q_{\text{l}}(z_{\text{l}}|x)} \right] \tag{17}$$

Using Jensen's equality, the ELBO can be derived as:

$$\begin{aligned} \text{ELBO} = {} & \mathbb{E}_{q_{\text{bio}}(z_{\text{bio}}|x)\, q_{\text{bat}}(z_{\text{bat}}|x)\, q_{\text{l}}(l|x)} [\log p(x \mid z_{\text{bio}}, z_{\text{bat},l})] \\ & - \text{KL}\big(q_{\text{bio}}(z_{\text{bio}} \mid x) \,||\, p(z_{\text{bio}})\big) \\ & - \text{KL}\big(q_{\text{bat}}(z_{\text{bat}} \mid x) \,||\, p(z_{\text{bat}})\big) \\ & - \text{KL}\big(q_{\text{l}}(l \mid x) \,||\, p(\text{l})\big). \end{aligned} \tag{18}$$

where the first term remains a reconstruction term, while the last three terms represent the KL divergence between the variational distributions and their corresponding prior distributions. Specifically, the prior for the library size $p(l)$ in the last term is modeled as a one-dimensional Gaussian, whereas both $p(z_{\text{bio}})$ and $p(z_{\text{batch}})$ are standard Gaussiansn. Notably, the key differences between the ELBO for scRNA-seq and that for IMC lie in the reconstruction term (introduced later) and the additonal scaling factor $z_{\text{l}}$ in scRNA-seq.

##### A.2. Zero-Inflated Negative Binomial (ZINB) Loss

To model the discrete and zero-inflated nature of scRNA-seq data, we adopt a Zero-Inflated Negative Binomial (ZINB) distribution as the form of data likelihood. Let $x_i \in \mathbb{N}_{\geq 0}$ denote the observed count for cell $i$. We denote by $\mu_i$ the mean expression rate, $r$ the dispersion parameter, and $\pi$ the zero-inflation probability. The ZINB mixture model is given by

$$P(X_i = x_i \mid \mu_i, r, \pi) = \pi\, \delta_0(x_i) + (1 - \pi)\, \text{NB}(x_i \mid r, \mu_i), \tag{19}$$

where

$$\text{NB}(x_i \mid r, \mu_i) = \binom{x_i + r - 1}{x_i} \left(\frac{r}{r + \mu_i}\right)^r \left(\frac{\mu_i}{r + \mu_i}\right)^{x_i}. \tag{20}$$

Then, the corresponding negative log-likelihood for a single observation $x_i$ can be derived as:

$$-\log P(X_i = x_i \mid \mu_i, r, \pi) = \begin{cases} -\log\Big[\pi + (1-\pi)\Big(\frac{r}{r+\mu_i}\Big)^r\Big] \\ \quad \text{if } x_i = 0 \\ \\ -\log(1-\pi) \; - \; \log\binom{x_i+r-1}{x_i} \\ \; - r \, \log\Big(\frac{r}{r+\mu_i}\Big) \\ \; - x_i \, \log\Big(\frac{\mu_i}{r+\mu_i}\Big) \\ \quad \text{if } x_i > 0 \end{cases} \tag{21}$$

Then summing over all cells $i = 1, \ldots, N$, the final ZINB loss can be obtained

$$\mathcal{L}_{\text{ZINB}} = \sum_{i=1}^{N} -\log\Big[\pi \, \delta_0(x_i) + (1-\pi) \, \text{NB}(x_i \mid r, \mu_i)\Big] \tag{22}$$

which can be explicitly expanded as:

$$\mathcal{L}_{\text{ZINB}} = \sum_{i:x_i=0} -\log\Big[\pi + (1-\pi)\Big(\frac{r}{r+\mu_i}\Big)^r\Big] + \sum_{i:x_i>0} \Big[-\log(1-\pi) - \log\binom{x_i+r-1}{x_i} - r \, \log\Big(\frac{r}{r+\mu_i}\Big) - x_i \, \log\Big(\frac{\mu_i}{r+\mu_i}\Big)\Big]$$

Consequently, the ELBO of scRNA-seq can be re-written using the negative $\mathcal{L}_{\text{ZINB}}$ as the reconstruction term instead of MSE in IMC data:

$$\begin{aligned} \text{ELBO} = & -\mathcal{L}_{\text{ZINB}} \\ & - \text{KL}\big(q_{\text{bio}}(z_{\text{bio}} \mid x) \,||\, p(z_{\text{bio}})\big) \\ & - \text{KL}\big(q_{\text{bat}}(z_{\text{bat}} \mid x) \,||\, p(z_{\text{bat}})\big) \\ & - \text{KL}\big(q_{\text{l}}(l \mid x) \,||\, p(\text{l})\big) \end{aligned} \tag{23}$$

#### A.3. Totoal loss function

In conclusion, similar to Eq. (9), by utilizing the ELBO from Eq. (23) as the backbone, incorporating adversarial disentanglement training introduced in the Eq. (4), Eq. (5) and Eq. (8), the total loss function of scRNA-seq is formulated as follows:

$$\mathcal{L}_{\text{total}} = -\text{ELBO} + \lambda_1 \mathcal{L}_{\text{ce}}\big(E_{\text{bio}}, D_{\text{bat}}\big) + \lambda_2 \mathcal{L}_{\text{ce}}\big(E_{\text{bat}}, C_{\text{bat}}\big) + \lambda_3 \mathcal{L}_{\text{orth}} \tag{24}$$

where $\lambda_1$, $\lambda_2$ and $\lambda_3$, are still weighting factors.

### B. Evaluation metrics

#### B.1. Bacth Effect Correction

##### ASW-batch (Average Silhouette Width by Batch)

This metric quantifies how well cells are mixed across batches. For each cell $i$ in batch $B_i$:

$$a(i) = \frac{1}{|B_i| - 1} \sum_{j \in B_i, \, j \neq i} \text{dist}(x_i, x_j),$$

$$b(i) = \min_{b \neq B_i} \left( \frac{1}{|b|} \sum_{j \in b} \text{dist}(x_i, x_j) \right).$$

The silhouette width for cell $i$ is

$$\text{SW}(i) = \frac{b(i) - a(i)}{\max\{a(i), b(i)\}},$$

and ASW-batch is the average of $\text{SW}(i)$ over all cells.

##### Graph Connectivity

A k-nearest neighbor graph in the low-dimensional space is construted firstly. Let $\mathcal{B}$ be the set of all batches. For each batch $b \in \mathcal{B}$, find the largest connected subgraph containing any cell from $b$, and let $|\mathcal{C}_b|$ be the number of cells from batch $b$ in that subgraph. The graph connectivity score is

$$\text{GC} = \frac{1}{|\mathcal{B}|} \sum_{b \in \mathcal{B}} \frac{|\mathcal{C}_b|}{|b|}.$$

##### iLISI (Integrated Local Inverse Simpson's Index)

For each cell $i$, consider its local neighborhood (e.g., k-nearest neighbors) and let $p_{i,b}$ be the fraction of neighbors belonging to batch $b$. The Inverse Simpson's Index (ISI) for cell $i$ is

$$\text{ISI}(i) = \frac{1}{\sum_{b \in \mathcal{B}} p_{i,b}^2}.$$

iLISI is then the average of ISI($i$) across all cells:

$$\text{iLISI} = \frac{1}{N}\sum_{i=1}^{N} \text{ISI}(i),$$

where $N$ is the total number of cells.

##### PCR (Principal Component Regression)

The integrated data is projected onto principal components and regress each principal component on batch labels (often using one-hot encoding). Let

$$R^2(\text{PC}_k \sim \text{batch})$$

be the coefficient of determination for the $k$-th principal component regressed on the batch. The **PCR** score is the average $R^2$ over the top $d$ principal components:

$$\text{PCR} = \frac{1}{d}\sum_{k=1}^{d} R^2(\text{PC}_k \sim \text{batch}).$$

Overall, These metrics have different optimal directions. For ASW-batch and PCR, lower values indicate better performance, while for Graph Connectivity and iLISI, higher values are better. To compute the average scores, we used the Python scib package. This package normalizes all metrics to a 0–1 scale, ensuring that higher scores consistently represent better performance across all four metrics. The average batch effect correction score used in this study is calculated as

$$\text{Average Score} = \frac{\text{ASW-batch} + \text{PCR} + \text{Graph Connectivity} + \text{iLISI}}{4}. \tag{25}$$

#### B.2. Biological conservation

##### ASW-cell (Average Silhouette Width by Cell Type)

Similar to ASW-batch but based on cell type rather than batch. For each cell $i$ with true cell type $T_i$:

$$a(i) = \frac{1}{|T_i| - 1}\sum_{j \in T_i, j \neq i} \text{dist}(x_i, x_j), \quad b(i) = \min_{t \neq T_i}\Big(\frac{1}{|t|}\sum_{j \in t} \text{dist}(x_i, x_j)\Big).$$

The silhouette width for cell $i$ is then

$$\text{SW}(i) = \frac{b(i) - a(i)}{\max\{a(i), b(i)\}},$$

and ASW-cell is the average SW($i$) across all cells.

##### ARI (Adjusted Rand Index)

ARI measures the agreement between true labels (cell types) and predicted cluster labels (e.g., via Leiden clustering). Let $n_{ij}$ be the number of elements shared by cluster $i$ and cell type $j$, and define

$$a_i = \sum_j n_{ij}, \quad b_j = \sum_i n_{ij}, \quad n = \sum_{i,j} n_{ij}.$$

Then ARI is given by

$$\text{ARI} = \frac{\sum_{i,j}\binom{n_{ij}}{2} - \Big[\sum_i \binom{a_i}{2}\sum_j \binom{b_j}{2}\Big]\Big/\binom{n}{2}}{\frac{1}{2}\Big[\sum_i \binom{a_i}{2} + \sum_j \binom{b_j}{2}\Big] - \Big[\sum_i \binom{a_i}{2}\sum_j \binom{b_j}{2}\Big]\Big/\binom{n}{2}}.$$

##### NMI (Normalized Mutual Information)

NMI also captures how well the clustering (predicted labels) matches the true cell types. Let $U$ be the set of true labels and $V$ be the set of predicted cluster labels. Define

$$\text{NMI}(U, V) = \frac{2\, I(U;V)}{H(U) + H(V)},$$

where $I(U;V)$ is the mutual information between $U$ and $V$, and $H(U)$ and $H(V)$ are their respective entropies.

In conclustion, similat to batch effect correction average score, we also use scib python package which maps all the evaluation values to 0-1 scale. Thus, the average biological score is calculated as

$$\text{Average Score} = \frac{\text{ASW-cell} + \text{ARI} + \text{NMI}}{3}. \tag{26}$$

#### B.3. Clustering performance

We use the Adjusted Rand Index (ARI), Normalized Mutual Information (NMI), and Accuracy (ACC) to evaluate the clustering performance. The definitions of ARI and NMI are provided in Section B.2. Here, ACC is defined as the optimal matching accuracy between the predicted cluster assignments and the ground truth labels.

Given a set of predicted clusters $\mathcal{C} = \{C_1, C_2, \ldots, C_k\}$ and the ground truth labels $\mathcal{L} = \{L_1, L_2, \ldots, L_k\}$, the accuracy is defined as the highest possible matching between these clusters and labels. Formally, the ACC is expressed as:

$$\text{ACC} = \frac{\max_{\pi \in \Pi} \sum_{i=1}^{k} \left|C_i \cap L_{\pi(i)}\right|}{n}$$

where:

- Π represents the set of all possible permutations of the cluster labels.
- $\pi$ is a specific permutation in Π.
- $|C_i \cap L_{\pi(i)}|$ denotes the number of data points common to the $i$-th predicted cluster and the $\pi(i)$-th ground truth label.
- $n$ is the total number of data points in the dataset.

#### C. Experiment results of IMC datasets

In this section, we present the evaluation results of our BioBatchNet and seven benchmark methods, including five batch effect correction metrics and three biological conservation metrics, assessed across three IMC datasets (refer to Tables S1, S2, and S3). Additionally, we showcase the UMAP embeddings generated by our BioBatchNet model alongside seven benchmark methods, with visualizations colored by batch and cell type, using the Damond (Figure S1) and HochSchulz (Figure S2) datasets for comparison.

##### C.1. Experiment results of IMMUcan_Cancer dataset

Table S1 presents the evaluation results for BioBatchNet and other seven benchmark methods across all metrics on the Immucan IMC dataset, including their average scores. Notably, our BioBatchNet achieves the highest average performance in both batch effect correction and biological conservation metrics.

**Table S1.** Comparison of batch effect correction and biological conservation scores across batch correction methods in IMMUcan_Cancer dataset.

| | Batch Effect Correction Score | | | | | Biological Conservation Score | | | |
|---|---|---|---|---|---|---|---|---|---|
| Model | iLISI | Graph connectivity | ASW-batch | PCR | Avg. score | ASW-cell | ARI | NMI | Avg. score |
| scVI | 0.21 | 0.99 | 0.92 | 0.49 | 0.65 | 0.56 | 0.24 | 0.44 | 0.41 |
| Harmony | 0.44 | 0.99 | 0.95 | 0.92 | 0.82 | 0.56 | 0.50 | 0.54 | 0.53 |
| BBKNN | 0.02 | 0.99 | 0.87 | 0.00 | 0.47 | 0.55 | 0.36 | 0.53 | 0.48 |
| Scanorama | 0.49 | 0.87 | 0.94 | 0.77 | 0.77 | 0.51 | 0.12 | 0.34 | 0.32 |
| Combat | 0.19 | 0.99 | 0.93 | 1.00 | 0.78 | 0.54 | 0.31 | 0.51 | 0.45 |
| iMAP | 0.07 | 0.99 | 0.90 | 0.90 | 0.72 | 0.51 | 0.19 | 0.37 | 0.36 |
| scDREAMER | 0.59 | 0.97 | 0.98 | 0.77 | 0.83 | 0.52 | 0.36 | 0.36 | 0.41 |
| **BioBatchNet** | 0.61 | 0.99 | 0.95 | 0.92 | 0.87 | 0.54 | 0.58 | 0.55 | 0.56 |

##### C.2. Experiment results of Damond_Pancreas dataset

Table S2 presents the evaluation results for BioBatchNet and other seven benchmark methods across all metrics on the Damond IMC dataset, including their average scores. Notably, our BioBatchNet achieves the second-highest performance in batch effect correction, while ranking first in both biological conservation and clustering performance. Figure S1 is the UMAP embedding of batch and biological information before and after batch effect correction across eight methods, including our BioBatchNet.

**Table S2.** Comparison of batch effect correction and biological conservation scores across batch correction methods in Damond_Pancreas dataset.

| | Batch effect correction Score | | | | | Biological conservation Score | | | |
|---|---|---|---|---|---|---|---|---|---|
| Model | iLISI | Graph connectivity | ASW-batch | PCR | Avg. score | ASW-cell | ARI | NMI | Avg. score |
| scVI | 0.34 | 0.96 | 0.87 | 0.69 | 0.71 | 0.56 | 0.56 | 0.54 | 0.56 |
| Harmony | 0.47 | 0.92 | 0.88 | 0.93 | 0.80 | 0.58 | 0.59 | 0.52 | 0.56 |
| BBKNN | 0.27 | 0.73 | 0.88 | 0.54 | 0.60 | 0.58 | 0.23 | 0.42 | 0.41 |
| Scanorama | 0.41 | 0.70 | 0.83 | 0.73 | 0.61 | 0.58 | 0.50 | 0.47 | 0.51 |
| Combat | 0.40 | 0.90 | 0.89 | 0.99 | 0.73 | 0.58 | 0.60 | 0.53 | 0.57 |
| iMAP | 0.00 | 0.65 | 0.71 | 0.07 | 0.48 | 0.54 | 0.05 | 0.28 | 0.29 |
| scDREAMER | 0.50 | 0.70 | 0.93 | 0.87 | 0.75 | 0.51 | 0.31 | 0.40 | 0.35 |
| **BioBatchNet** | 0.48 | 0.88 | 0.88 | 0.81 | 0.77 | 0.57 | 0.58 | 0.54 | 0.56 |

##### C.3. Experiment results of Hoch_Melanoma dataset

Table S3 presents the evaluation results for BioBatchNet and other seven benchmark methods across all metrics on the HochSchulz IMC dataset, including their average scores. Notably, our BioBatchNet achieves the second-highest performance in batch effect correction, while ranking first in both biological conservation and clustering performance. Figure S2 is the UMAP embedding of batch and biological information before and after batch effect correction across eight methods, including our BioBatchNet.

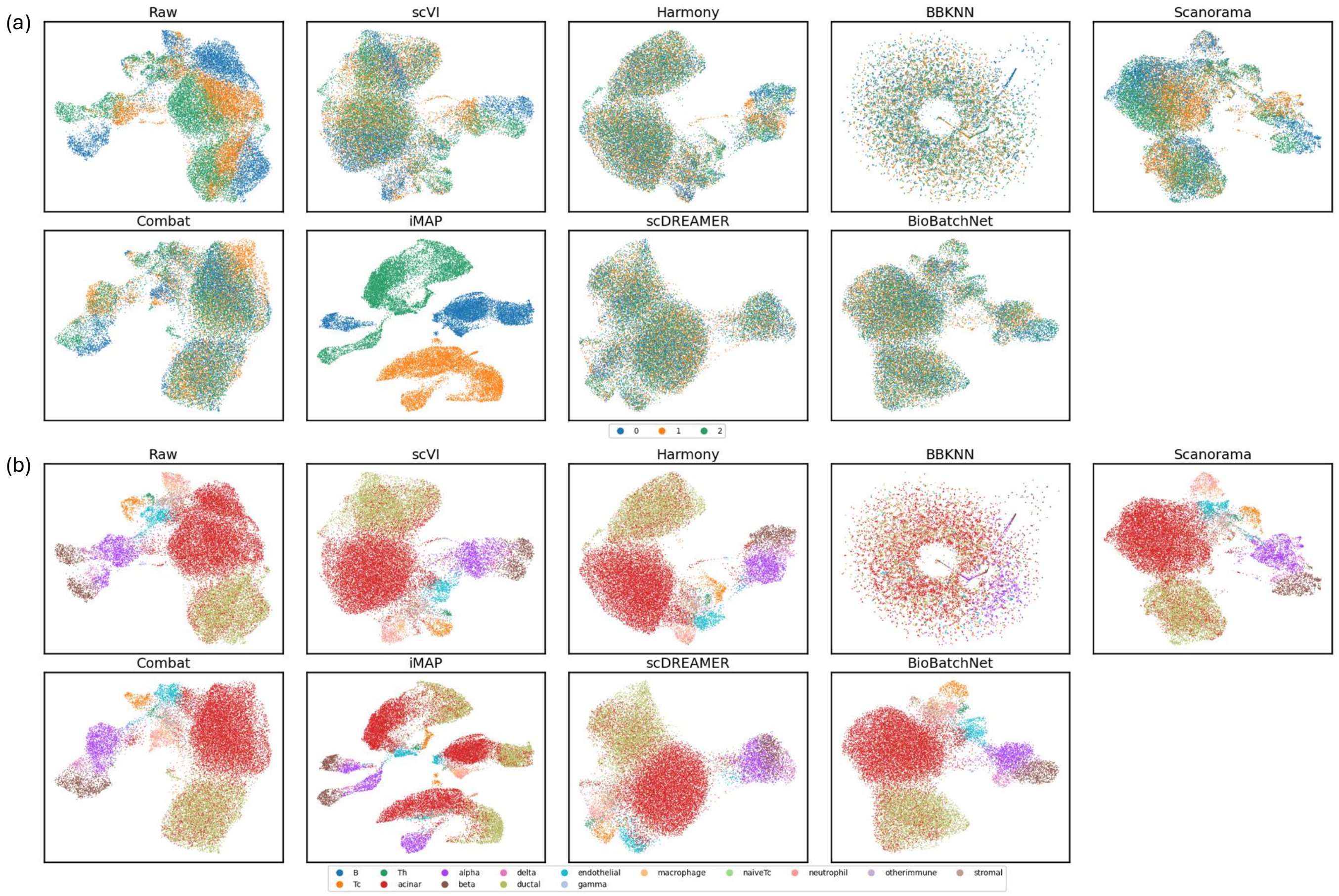


Fig. S1: Comparison of methods for Damond_Pancreas dataset. **(a)** UMAP visualization illustrating of batch effect correction in latent biological representations. **(b)** UMAP visualization of biological signal conservation in latent biological representations.

**Table S3.** Comparison of batch effect correction and biological conservation scores across batch correction methods in Hoch_Melanoma dataset.

| | Batch effect correction | | | | | Biological Conservation Score | | | |
|---|---|---|---|---|---|---|---|---|---|
| **Model** | **iLISI** | **Graph connectivity** | **ASW-batch** | **PCR** | **Avg. score** | **ASW-cell** | **ARI** | **NMI** | **Avg. score** |
| scVI | 0.09 | 0.98 | 0.84 | 0.23 | 0.53 | 0.53 | 0.26 | 0.38 | 0.39 |
| Harmony | 0.15 | 0.98 | 0.83 | 0.71 | 0.67 | 0.56 | 0.47 | 0.55 | 0.53 |
| BBKNN | 0.05 | 0.98 | 0.88 | 0.00 | 0.48 | 0.55 | 0.46 | 0.53 | 0.51 |
| Scanorama | 0.17 | 0.54 | 0.80 | 0.93 | 0.61 | 0.45 | 0.09 | 0.08 | 0.21 |
| Combat | 0.12 | 0.97 | 0.90 | 1.00 | 0.75 | 0.55 | 0.42 | 0.48 | 0.48 |
| iMAP | 0.00 | 0.90 | 0.82 | 0.21 | 0.48 | 0.49 | 0.04 | 0.18 | 0.24 |
| scDREAMER | 0.17 | 0.94 | 0.86 | 0.86 | 0.71 | 0.52 | 0.25 | 0.29 | 0.35 |
| **BioBatchNet** | 0.15 | 0.98 | 0.85 | 0.78 | 0.69 | 0.55 | 0.65 | 0.54 | 0.58 |

#### D. Experiment results of scRNA-seq datasets

In this section, we present the evaluation results of our BioBatchNet model alongside eight benchmark methods, focusing on batch effect correction and biological conservation across four scRNA-seq datasets (see Tables S6, S4, S5, and S7). Moreover, we also show the UAMP embedding of our BioBatchNet and eight benchmark methods across four scRNA-seq datasets in the Figure S5, S3, S4 and S6.

##### D.1. Experiment results of Human pancreas scRNA-seq data

Table S4 summarizes the evaluation results for BioBatchNet alongside eight benchmark methods across all metrics on the Human pancreas scRNA-seq dataset, including their average scores. Notably, BioBatchNet still demonstrates the best performance in batch effect correction, ranks second in biological conservation and clustering performance overall. Figure S3 illustrates the UMAP embeddings of batch and biological information before and after batch effect correction for BioBatchNet and the eight benchmark methods.

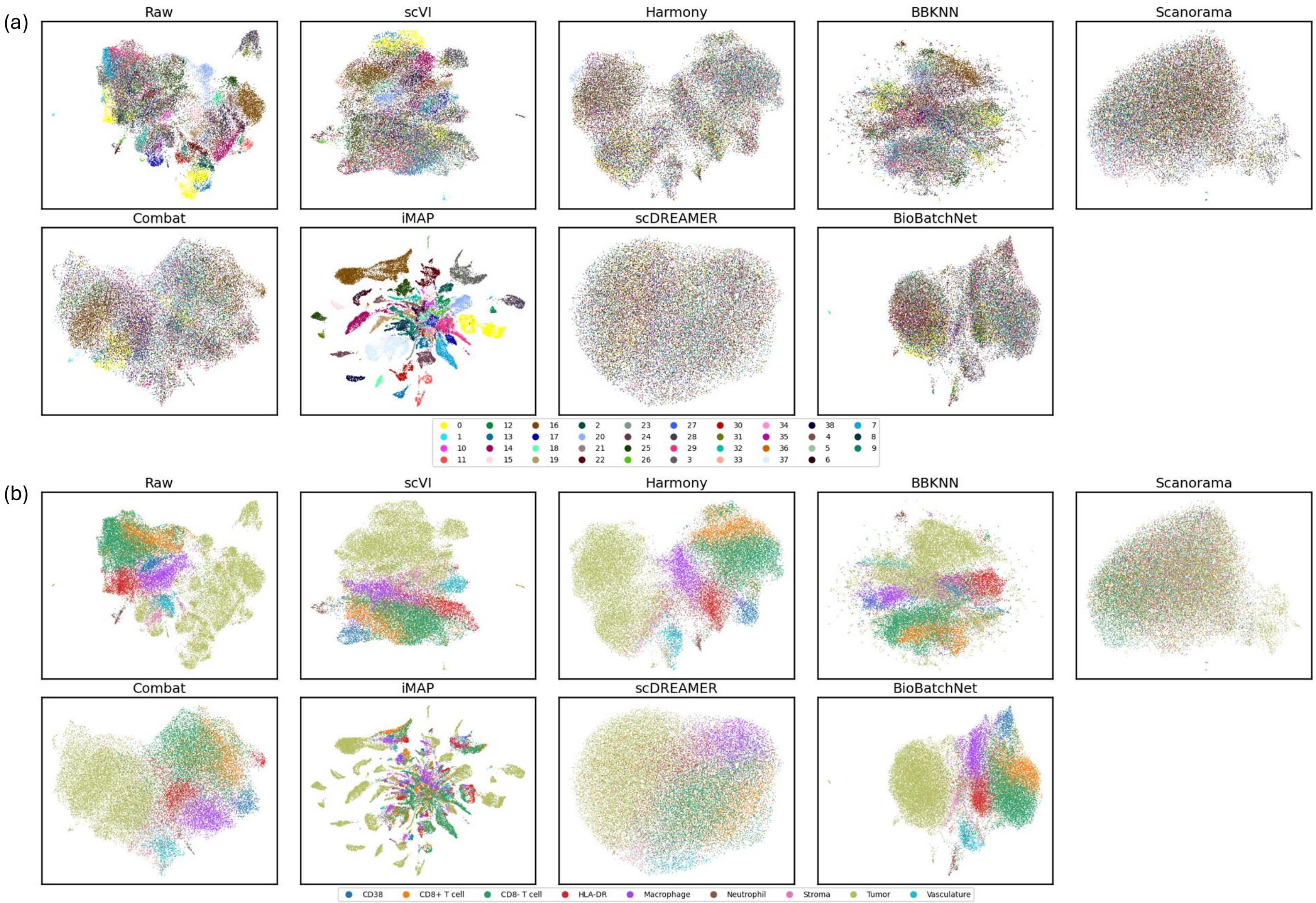


Fig. S2: UMAP plots of Hoch_Melanoma dataset. **(a)** is the visualization of batch information before and after batch effect correction across eight methods, including our BioBatchNet, using the HochSchulz IMC dataset. Different colors represent distinct batches. **(b)** is the visualization of biological information before and after integration, with cells color-coded according to their respective cell types.

**Table S4.** Comparison of batch effect correction and biological conservation scores across batch correction methods in Human pancreas dataset.

| | Batch effect correction | | | | | Biological Conservation Score | | | |
|---|---|---|---|---|---|---|---|---|---|
| **Model** | **iLISI** | **Graph connectivity** | **ASW-batch** | **PCR** | **Avg. score** | **ASW-cell** | **ARI** | **NMI** | **Avg. score** |
| scVI | 0.26 | 0.98 | 0.88 | 0.54 | 0.67 | 0.62 | 0.95 | 0.92 | 0.83 |
| Harmony | 0.35 | 0.96 | 0.90 | 0.83 | 0.76 | 0.65 | 0.95 | 0.91 | 0.84 |
| BBKNN | 0.01 | 0.98 | 0.84 | 0.00 | 0.46 | 0.59 | 0.95 | 0.91 | 0.82 |
| Scanorama | 0.28 | 0.86 | 0.82 | 0.55 | 0.63 | 0.59 | 0.28 | 0.61 | 0.49 |
| Combat | 0.17 | 0.97 | 0.92 | 1.00 | 0.77 | 0.62 | 0.84 | 0.86 | 0.77 |
| iMAP | 0.44 | 0.87 | 0.88 | 0.85 | 0.76 | 0.62 | 0.92 | 0.86 | 0.80 |
| scDREAMER | 0.42 | 0.99 | 0.87 | 0.85 | 0.78 | 0.72 | 0.96 | 0.93 | 0.87 |
| scDML | 0.42 | 0.93 | 0.88 | 0.89 | 0.78 | 0.77 | 0.94 | 0.92 | 0.88 |
| **BioBatchNet** | 0.48 | 0.99 | 0.90 | 0.91 | 0.83 | 0.61 | 0.94 | 0.91 | 0.83 |

#### D.2. Experiment results of Human immune scRNA-seq data

Table S5 presents the evaluation results for BioBatchNet and eight benchmark methods across all metrics on the Human immune scRNA-seq dataset, including their average scores. Notably, BioBatchNet secures second place in both batch effect correction and biological conservation simultaneously, still demonstrating strong competitive performance. Figure S4 illustrates the UMAP embeddings of batch and biological information before and after batch effect correction for BioBatchNet and the eight benchmark methods.

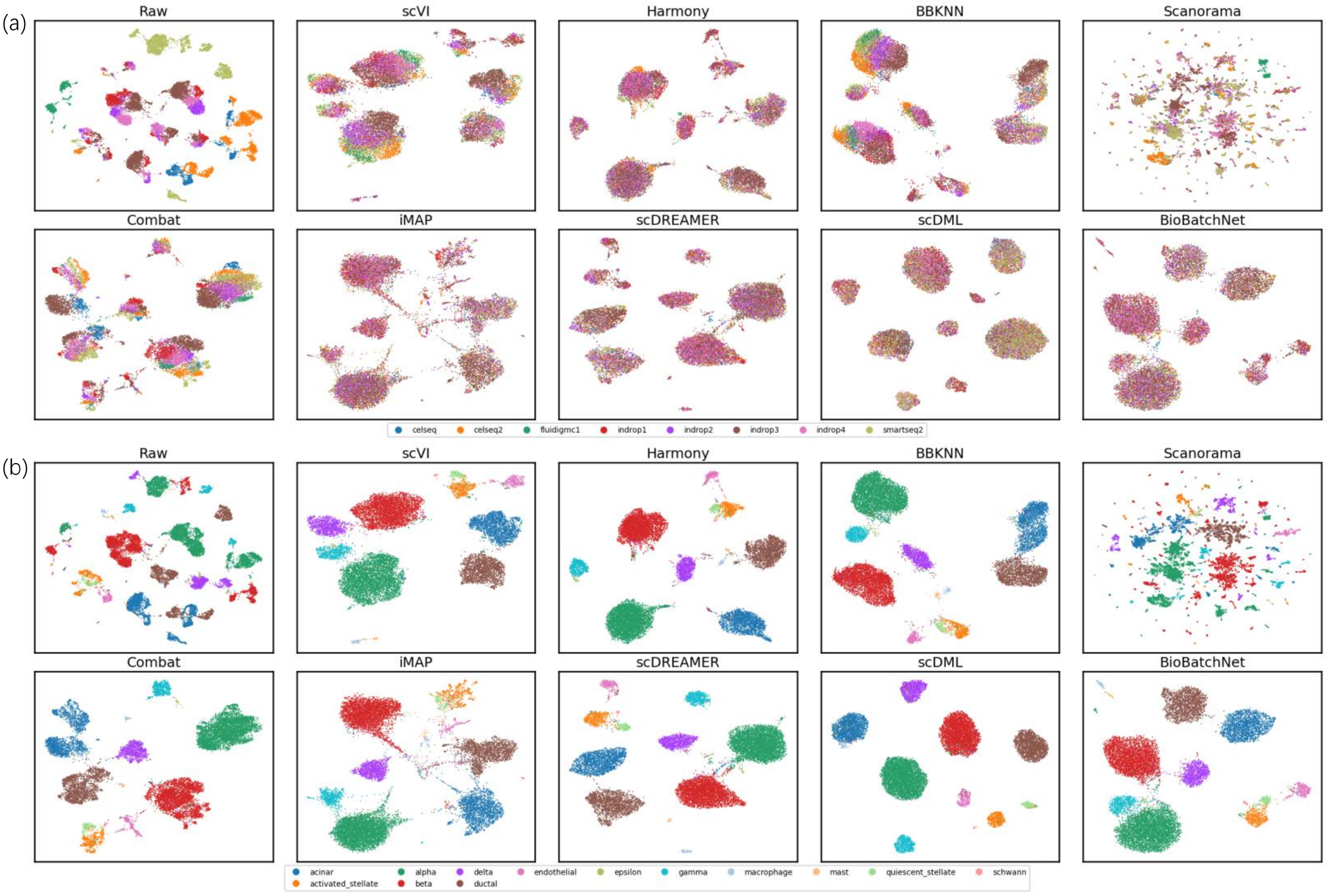


Fig. S3: Comparison of methods for Human pancreas dataset. **(a)** UMAP visualization illustrating of batch effect correction in latent biological representations. **(b)** UMAP visualization of biological signal conservation in latent biological representations.

**Table S5.** Comparison of batch effect correction and biological conservation scores across batch correction methods in Human immune dataset.

| | Batch effect correction | | | | | Biological Conservation Score | | | |
|---|---|---|---|---|---|---|---|---|---|
| **Model** | **iLISI** | **Graph connectivity** | **ASW-batch** | **PCR** | **Avg. score** | **ASW-cell** | **ARI** | **NMI** | **Avg. score** |
| scVI | 0.30 | 0.97 | 0.90 | 0.67 | 0.71 | 0.55 | 0.50 | 0.72 | 0.59 |
| Harmony | 0.29 | 0.96 | 0.94 | 0.80 | 0.75 | 0.54 | 0.73 | 0.76 | 0.68 |
| BBKNN | 0.13 | 0.99 | 0.88 | 0.00 | 0.50 | 0.54 | 0.75 | 0.80 | 0.70 |
| Scanorama | 0.22 | 0.83 | 0.86 | 0.31 | 0.56 | 0.54 | 0.27 | 0.52 | 0.44 |
| Combat | 0.29 | 0.97 | 0.93 | 1.00 | 0.80 | 0.54 | 0.73 | 0.78 | 0.68 |
| iMAP | 0.34 | 0.94 | 0.93 | 0.74 | 0.74 | 0.52 | 0.60 | 0.67 | 0.60 |
| scDREAMER | 0.28 | 0.94 | 0.86 | 0.59 | 0.67 | 0.61 | 0.60 | 0.77 | 0.66 |
| scDML | 0.28 | 0.86 | 0.86 | 0.86 | 0.72 | 0.52 | 0.44 | 0.60 | 0.52 |
| **BioBatchNet** | 0.31 | 0.93 | 0.91 | 0.71 | 0.72 | 0.56 | 0.74 | 0.78 | 0.69 |

#### D.3. Experiment results of Macaque retina dataset

Table S6 summarizes the evaluation results for BioBatchNet alongside eight benchmark methods across all metrics on the Macaque retina scRNA-seq dataset, including their average scores. Notably, BioBatchNet demonstrates the best performance in batch effect correction, ranks second in biological conservation, and achieves the highest clustering performance overall. Figure S5 illustrates the UMAP embeddings of batch and biological information before and after batch effect correction for BioBatchNet and the eight benchmark methods.

#### D.4. Experiment results of Mouse brain scRNA-seq data

Table S7 presents the evaluation results for BioBatchNet and eight benchmark methods across all metrics on the Mouse brain scRNA-seq dataset, including their average scores. BioBatchNet demonstrates superior performance in both batch effect correction and biological conservation, excelling simultaneously in these key areas. Notably, BioBatchNet achieves significantly higher clustering performance, effectively overcoming

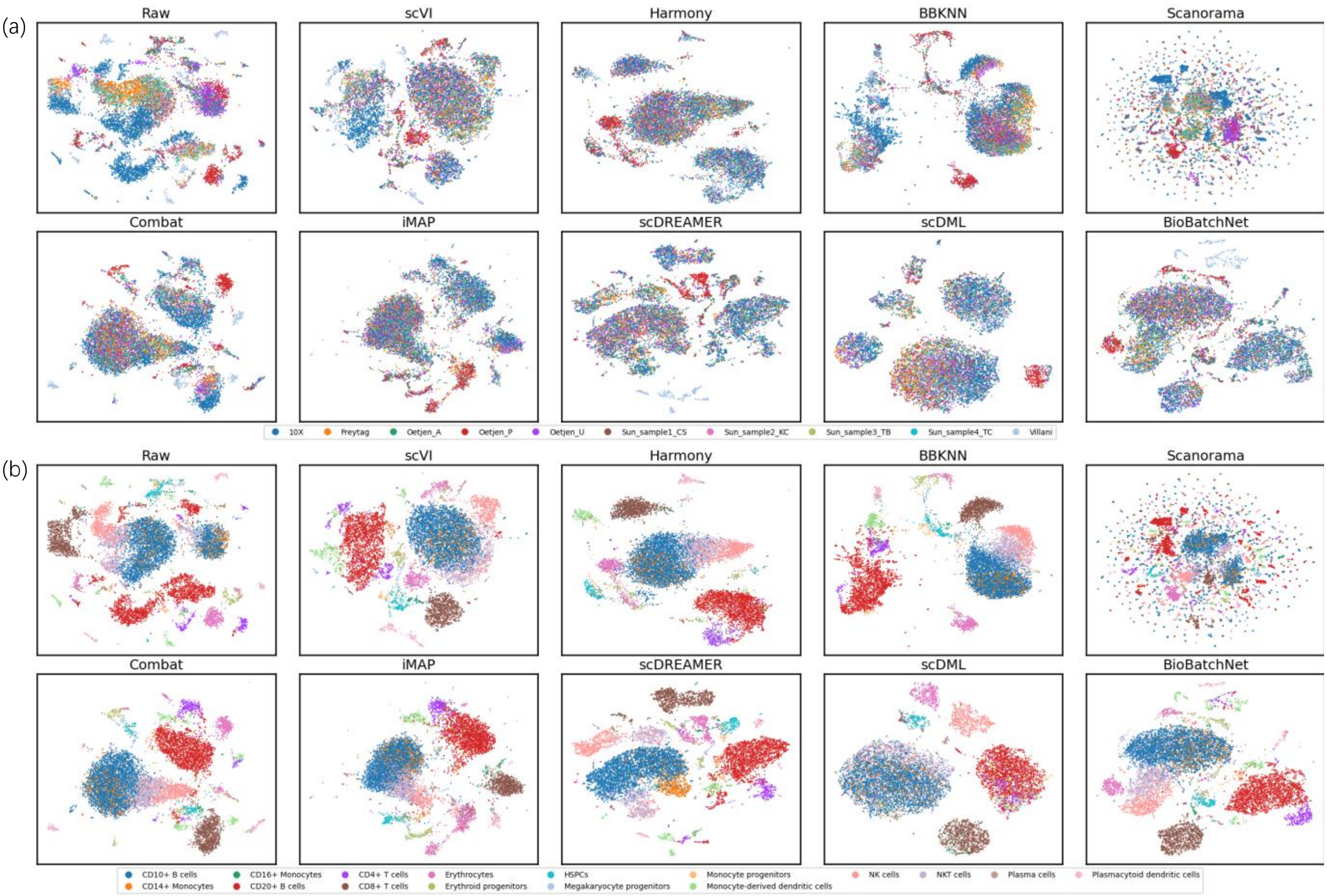


Fig. S4: Comparison of methods for Human immune dataset. **(a)** UMAP visualization illustrating of batch effect correction in latent biological representations. **(b)** UMAP visualization of biological signal conservation in latent biological representations.

**Table S6.** Comparison of batch effect correction and biological conservation scores across batch correction methods in Macaque retina dataset.

| | Batch effect correction | | | | | Biological Conservation Score | | | |
|---|---|---|---|---|---|---|---|---|---|
| **Model** | **iLISI** | **Graph connectivity** | **ASW-batch** | **PCR** | **Avg. score** | **ASW-cell** | **ARI** | **NMI** | **Avg. score** |
| scVI | 0.34 | 0.99 | 0.92 | 0.83 | 0.77 | 0.60 | 0.87 | 0.90 | 0.79 |
| Harmony | 0.52 | 0.98 | 0.94 | 0.90 | 0.84 | 0.56 | 0.74 | 0.80 | 0.70 |
| BBKNN | 0.01 | 0.99 | 0.87 | 0.00 | 0.47 | 0.54 | 0.89 | 0.89 | 0.77 |
| Scanorama | 0.29 | 0.75 | 0.87 | 0.28 | 0.55 | 0.55 | 0.52 | 0.71 | 0.59 |
| Combat | 0.22 | 0.99 | 0.95 | 1.00 | 0.79 | 0.55 | 0.70 | 0.76 | 0.67 |
| iMAP | 0.38 | 0.97 | 0.90 | 0.49 | 0.69 | 0.54 | 0.48 | 0.66 | 0.56 |
| scDREAMER | 0.53 | 1.00 | 0.88 | 0.81 | 0.81 | 0.68 | 0.79 | 0.87 | 0.78 |
| scDML | 0.53 | 0.95 | 0.93 | 0.92 | 0.83 | 0.79 | 0.94 | 0.93 | 0.89 |
| **BioBatchNet** | 0.62 | 0.98 | 0.97 | 0.96 | 0.88 | 0.57 | 0.96 | 0.95 | 0.83 |

noise and delivering robust results. Figure S6 illustrates the UMAP embeddings of batch and biological information before and after batch effect correction for BioBatchNet and the eight benchmark methods.

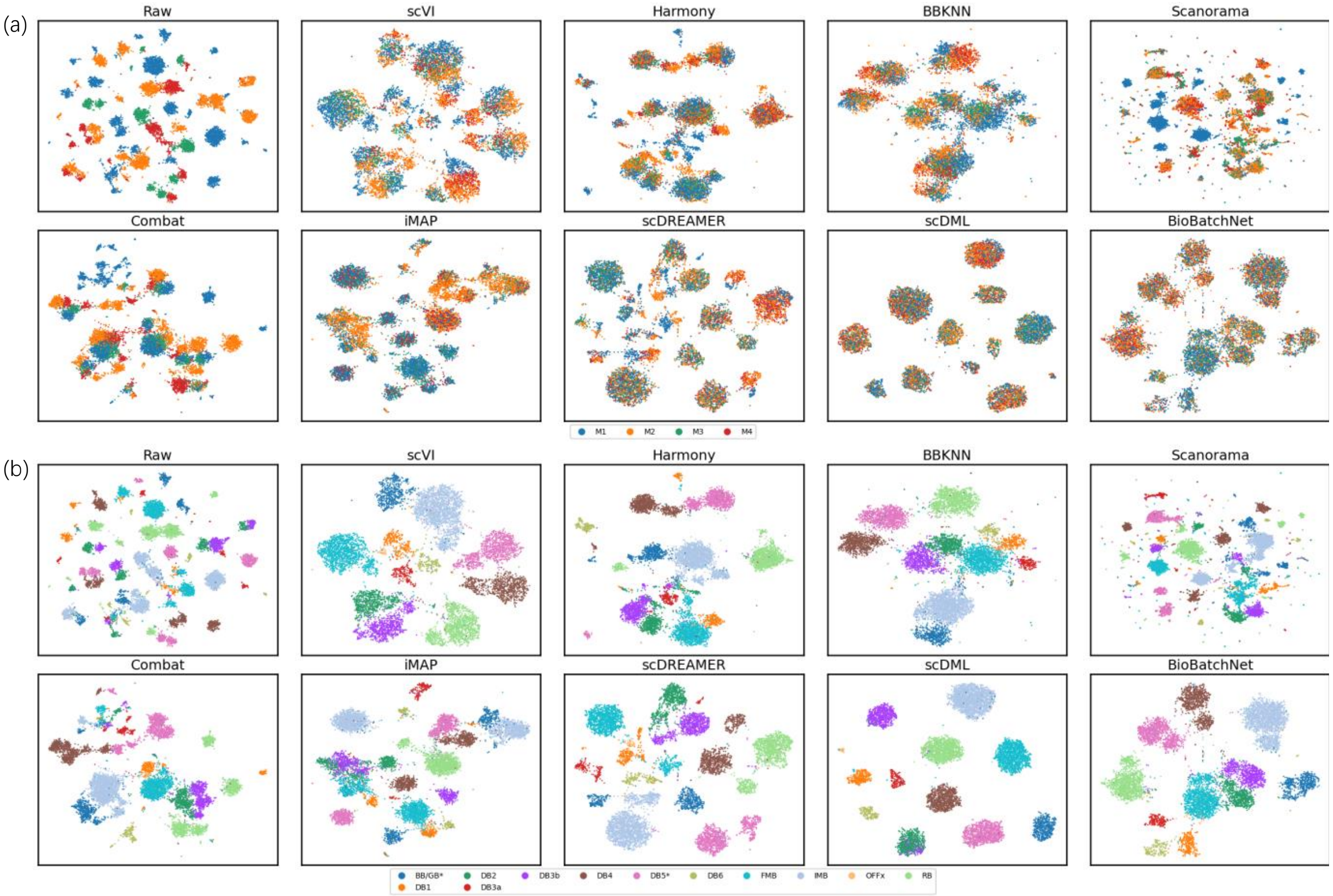


Fig. S5: Comparison of methods for Macaque retina dataset. **(a)** UMAP visualization illustrating of batch effect correction in latent biological representations. **(b)** UMAP visualization of biological signal conservation in latent biological representations.

**Table S7.** Evaluation results of BioBatchNet compared to other batch correction methods using mouse brain dataset.

| | Batch effect correction | | | | | Biological Conservation Score | | | |
|---|---|---|---|---|---|---|---|---|---|
| **Model** | **iLISI** | **Graph connectivity** | **ASW-batch** | **PCR** | **Avg. score** | **ASW-cell** | **ARI** | **NMI** | **Avg. score** |
| scVI | 0.04 | 0.93 | 0.82 | 0.90 | 0.67 | 0.56 | 0.28 | 0.59 | 0.48 |
| Harmony | 0.00 | 0.96 | 0.85 | 0.72 | 0.63 | 0.57 | 0.32 | 0.59 | 0.49 |
| BBKNN | 0.00 | 0.88 | 0.62 | 0.00 | 0.38 | 0.55 | 0.01 | 0.25 | 0.40 |
| Scanorama | 0.00 | 0.90 | 0.70 | 0.16 | 0.44 | 0.56 | 0.26 | 0.57 | 0.47 |
| Combat | 0.00 | 0.95 | 0.75 | 1.00 | 0.68 | 0.59 | 0.28 | 0.56 | 0.48 |
| iMAP | 0.17 | 0.87 | 0.81 | 0.94 | 0.70 | 0.57 | 0.30 | 0.58 | 0.49 |
| scDREAMER | 0.13 | 0.92 | 0.80 | 0.94 | 0.70 | 0.67 | 0.29 | 0.58 | 0.51 |
| scDML | 0.19 | 0.81 | 0.78 | 0.90 | 0.67 | 0.57 | 0.65 | 0.66 | 0.63 |
| **BioBatchNet** | 0.31 | 0.89 | 0.94 | 0.99 | 0.78 | 0.52 | 0.86 | 0.71 | 0.71 |

#### E. Clustering performance

##### E.1. Result of constrained pairwise clustering method

Notably, in the Damond dataset, acinar and ductal cells are subtypes of exocrine cells. However, due to the limited information available from IMC data, some acinar cells may be misclassified as ductal cells. Such incorrect labeling can lead to unreliable priors and suboptimal clustering results. To address this issue and ensure the accuracy of prior knowledge, we combine ductal and acinar cells into a single exocrine category. Figure S7 demonstrate the visulization results of Damond and HochSchulz IMC datasets after CPC. It can be seen that the CPC successfully overcomes the imbalanced and overlapping issue in IMC data, showing the superior clustering performance.

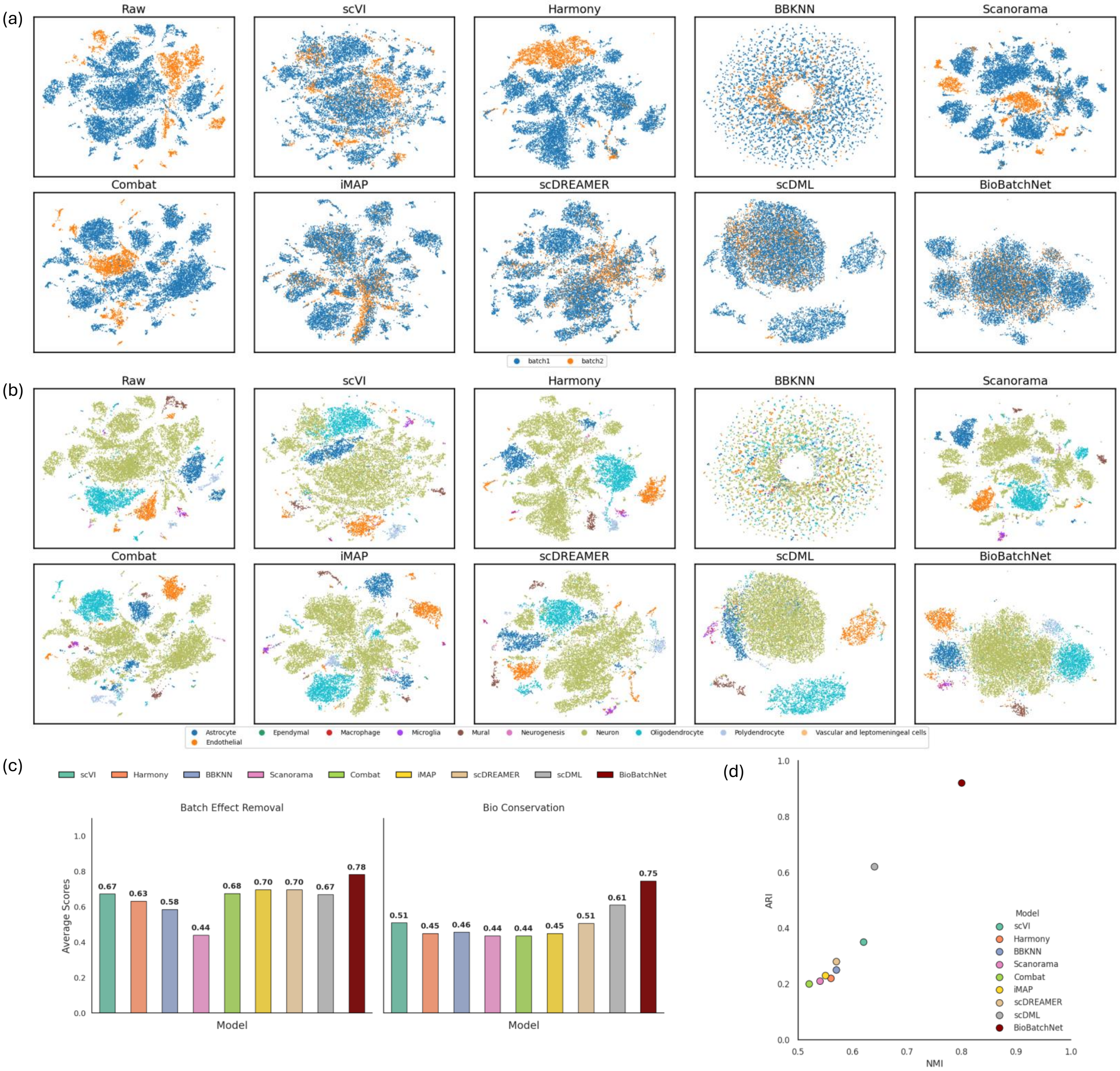


Fig. S6: Comparison of methods for Mouse brain dataset. **(a)** UMAP visualization illustrating of batch effect correction in latent biological representations. **(b)** UMAP visualization of biological signal conservation in latent biological representations. **(c)** UMAP visualization of batch information encoded in latent batch-specific representations. **(d)** Quantitative comparison of batch effect correction and biological conservation scores.

#### E.2. Limitation of deep embedded clustering methods

In this study, we highlight the disadvantages of deep embedded clustering (DEC) methods, which ultimately led us to exclude it from our CPC. DEC struggles to handle imbalanced and overlapping data effectively. As illustrated in Figure S8, tumor cells in (a) and (c), as well as exocrine cells in (b), are separated incorrectly, resulting in smaller clusters being mistakenly merged. This limitation arises because DEC tends to produce uniform clusters, rendering it unsuitable for the IMC dataset. The figure underscores DEC's inability to address the unique challenges of IMC data, particularly its imbalance and overlap.

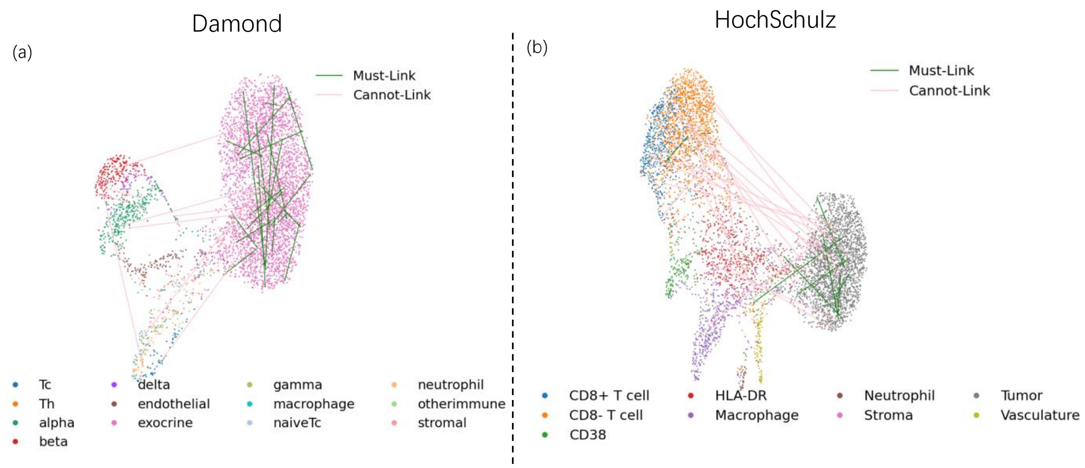


Fig. S7: CPC results for Damond and HochSchulz IMC data

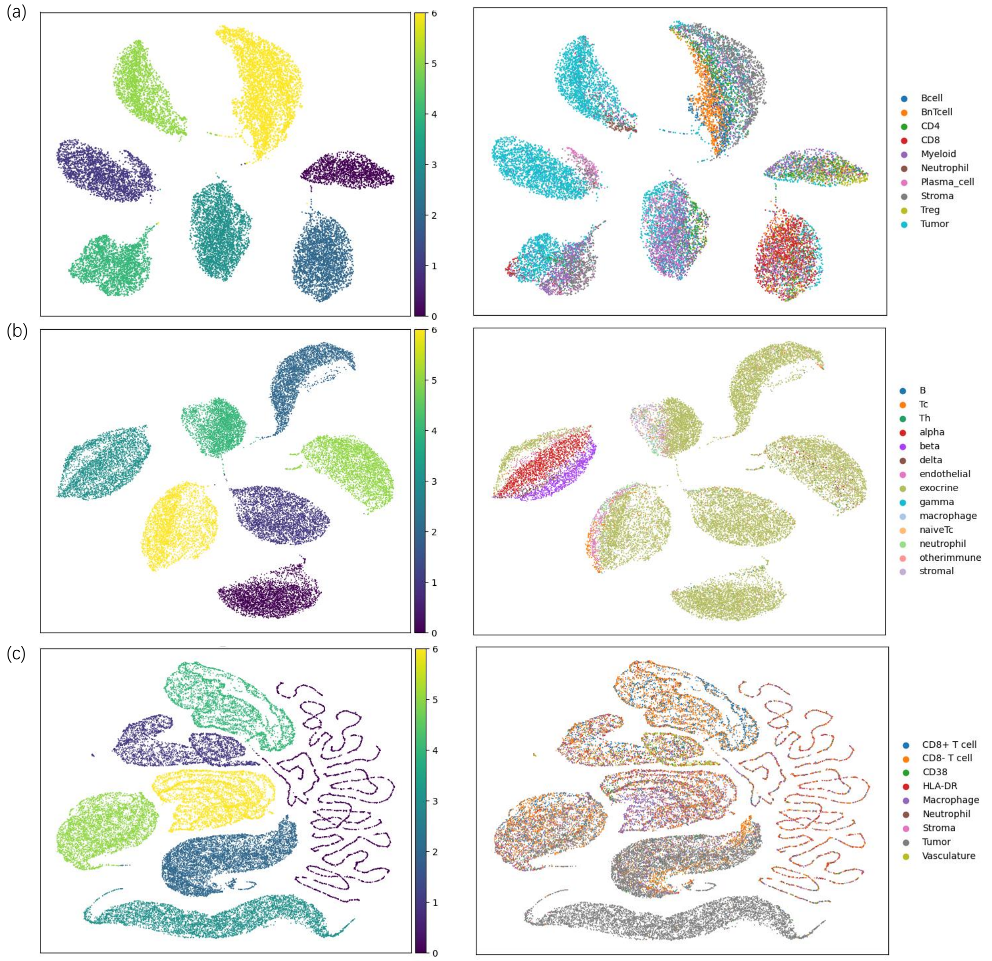


Fig. S8: DEC results across three IMC datasets. **(a)**, **(b)**, **(c)** are the IMMUcan, Damond, HochSchulz IMC dataset respectively.
